# A knowledge-guided approach to recovering important rare signals from high-dimensional single-cell data

**DOI:** 10.1101/2025.10.01.679835

**Authors:** Zhenghao Zhang, Jiamin Chen, Haoran Wu, Kelly Yichen Li, Peter D. Adams, Pamela Itkin-Ansari, Kevin Y. Yip

**Author notes:** These authors contributed equally: Zhenghao Zhang, Jiamin Chen, Haoran Wu.

## Abstract

Single-cell transcriptomic data are high-dimensional, with many genes profiled in each cell. Dimensionality reduction is routinely applied to improve interpretability, remove noise and redundancy, and enable visualization. Most existing methods aim at preserving the most prominent data properties, which can lead to omission of rare but important signals. Here we propose a novel framework that uses knowledge-derived genes of interest to guide dimensionality reduction, which can help cluster rare cells and separate highly similar cell sub-populations. We demonstrate the utility of our framework in identifying endocrine cell subtypes in the pancreatic islet, highly similar hematopoietic sub-populations, and rare senescent cells.

## Background

In single-cell transcriptomics, each cell can be considered as a point in a high-dimensional space formed by the usually thousands of genes profiled. The high-dimensional nature makes single-cell transcriptomic data difficult to interpret and visualize. As a result, dimensionality reduction methods are typically applied to find a low-dimensional representation of the data (called an “embedding”) that preserves properties of the original data. In addition to improving interpretability, the embedding also reduces noise and redundancy. A variety of downstream analysis tasks take the embedding as input, such as cell clustering, cell type annotation, and trajectory inference. Given its importance, many methods have been proposed for performing dimensionality reduction on single-cell transcriptomic data ^1–13^, which have been extensively compared in benchmarking studies ^14–16^.

How well the original data are preserved in the embedding can be quantified in different ways, such as the amount of variance explained, preservation of cell-cell distances or local cell neighborhoods, and the extent to which the original data can be reconstructed. If the embedding has a small number of dimensions, it is inevitable that some information would be lost, and typically dimensionality reduction methods resort to preserving the most prominent signals and sacrificing other signals. This can create the issue that rare but biologically interesting signals are not preserved. For example, if only a small proportion of cells belong to a cell type, information about these cells may be lost in the embedding since the dimensionality reduction method considers it more important to preserve information about the majority of the cells. As a result, these rare cells may not be clustered together in the embedding, which in turn makes it difficult to identify them from the data. As another example, if a parent cell type contains two subtypes but cells in these subtypes express most genes at similar levels except for a small number of subtype-specific genes, the embedding may capture more information about the commonly expressed genes as it separates the parent cell type from other cell types. Consequently, subtype-specific information could be lost and the two subtypes are not separated from each other well in the embedding.

A useful way to deal with this issue is to use some external knowledge to guide the dimensionality reduction process, such that extra attention is given to signals that are considered important according to the external knowledge. Currently, there are a few semi-supervised dimensionality reduction methods that adopt this idea, including MCML ^17^, netAE ^18^, and scnym ^19^. All of them use examples of cells of certain populations as external knowledge. Supplying these example cells correctly requires substantial expertise. If these example cells are curated manually, the process could be tedious, time-consuming, and error-prone.

Here we propose a novel approach to performing knowledge-guided dimensionality reduction of single-cell transcriptomic data, which takes a list of genes of interest as knowledge input, such as marker genes of a rare cell type. The embedding will be instructed to preserve more information about these genes, which in turn helps identify cells of this rare cell type as well as additional novel marker genes of it. The marker genes in the knowledge input can also help separate highly similar subpopulations and retain information about other genes that express differently between them, which represent another type of rare signals. As compared to example cells, marker genes can be more easily obtained either from domain knowledge or differential expression analysis of previous single-cell transcriptomic data. We give three demonstrations of the use of our method, in identifying different subtypes of pancreatic islet cells, separating highly similar hematopoietic sub-populations, and detecting rare senescent cells, respectively. We show that our method can tolerate noisy and incomplete knowledge inputs, and does not create data distortion problems.

## Results

### SAKURA: A knowledge-guided dimensionality reduction framework for single-cell transcriptomic data

In order to use external knowledge to guide the cell embedding process, we developed a general framework called <u>S</u>ingle-cell data <u>A</u>nalysis with <u>K</u>nowledge inputs from <u>U</u>ser using <u>R</u>egularized <u>A</u>utoencoders (SAKURA). The backbone of SAKURA is an encoder-decoder network that forms an embedding of the cells in its bottleneck layer with an aim to maximally reconstruct the input data (Figure 1a, Methods). Various modules can be flexibly attached to the backbone to achieve knowledge-guided embedding (Figure 1a). For instance, similar to previous methods ^17–19^, if example cells of some cell types are available, this knowledge can be supplied through the “Cell Labels” module to improve clustering of cells in the embedding by their cell types.

**Fig. 1:**
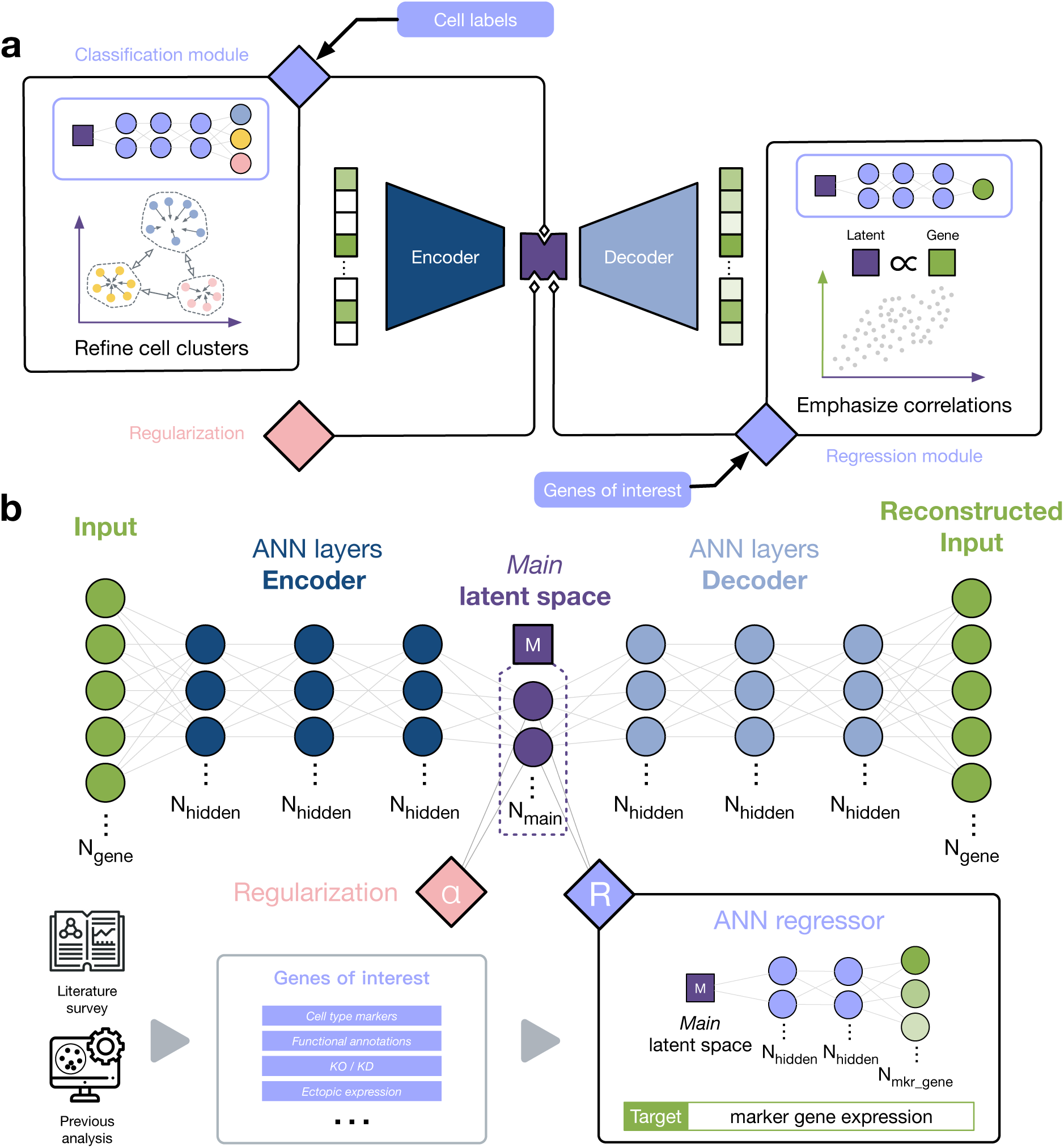
The SAKURA framework and its knowledge modules. **a,** The encoder-decoder backbone of SAKURA and three example modules that can be attached to it. Each module takes a different type of knowledge input to guide the cell embedding process. **b,** The Genes of Interest module that takes a list of genes as knowledge input and encourages the cell embedding to retain information about their expression in individual cells using regression.

Yet importantly, SAKURA can also take other types of knowledge inputs through other modules. As the focus of the current study, the “Genes of Interest” (GOI) module takes a list of genes as external knowledge input, for which the embedding is asked to retain their expression information (Figure 1b, Methods). These genes can be selected for different reasons. They can be marker genes of cell types of interest, especially rare cell types that have few cells in the data set. Alternatively, the genes can be those that have a functional annotation of interest, such as genes that belong to a biological pathway. The genes can also be those that have been experimentally perturbed, such as the ones knocked out (KO), knocked down (KD), or ectopically expressed in a specific cell type. In all cases, information about the expression of these genes is retained in the embedding by attaching a regression module to the bottleneck layer (Figure 1b), which uses the cell embedding to infer expression levels of the genes of interest. If the information is well-retained, expression levels of the genes can be accurately inferred by the embedding. On the other hand, if the information is not well-retained, regression accuracy would be low, and correspondingly a penalty will be added to the loss function to inform the model training process that the embedding is not preferred. The resulting embedding produced by SAKURA maintains a balance between preserving general properties of the data and specific information about the genes of interest (Methods).

### Improved identification and characterization of pancreatic islet endocrine cells by knowledge-guided analysis

As the first demonstration of SAKURA, we used it to identify endocrine cells in the pancreatic islet during fetal development. Endocrine cells within the human pancreatic islet can be classified into five major subtypes based on the hormones they secrete, namely glucagon (GCG)-secreting *α* cells, insulin (INS)-secreting *β* cells, somatostatin (SST)-secreting *δ* cells, ghrelin (GHRL)-secreting *ε* cells, and pancreatic polypeptide (PPY)-secreting *γ* cells (also called PP cells) ^20–26^. To determine whether these cell types are distinguishable during fetal development, we downloaded scRNA-seq data produced from pancreas samples from a published human fetal cell atlas ^27^. We first processed the data using a standard unsupervised analysis pipeline (PCA followed by clustering; Methods). In the cell clusters formed, the five hormone genes are primarily expressed in islet endocrine cells according to the annotations provided by the original authors but the boundaries between the five subtypes of islet endocrine cells are unclear (Figure S1). Even when we extracted these cells and re-clustered them without the presence of the other cell types (acinar cells, ductal cells, etc.), the expression patterns of the five genes are still fairly similar in the resulting clusters (Figure 2a), making it difficult to identify cells of each islet endocrine subtype.

**Fig. 2:**
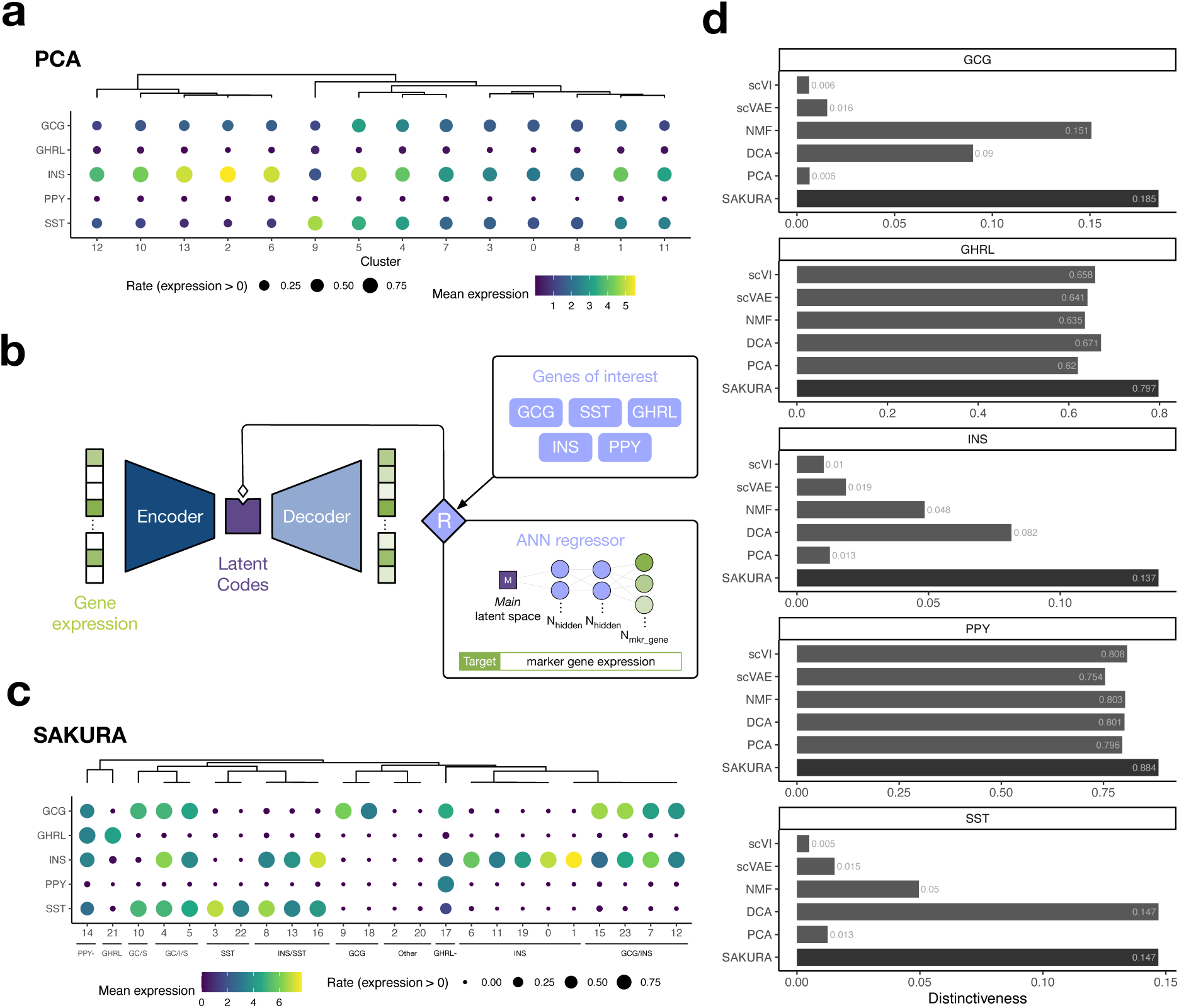
Identifying islet cell clusters by the knowledge-guided analysis of SAKURA. **a,** Expression of the five hormone genes in the clusters of the extracted cells identified by the unsupervised pipeline. **b,** The Genes of Interest module added to SAKURA that helps separate cells with different expression levels of the five hormone genes. **c,** Expression of the five hormone genes in the cell clusters produced from SAKURA’s embedding of the extracted cells. In Panels a and c, a complete-link hierarchical clustering of the cell clusters was performed based on cosine similarity of the detection rates of the five hormone genes. Clusters with similar expression patterns of the five genes are grouped together and given a common descriptive label in Panel c. **d,** Distinctiveness of the expression of the hormone genes in cell clusters produced from SAKURA’s and five other embedding methods’ embeddings. For each gene, the distinctiveness shown is the average across all clusters.

The poor separation of cells from the five islet endocrine subtypes indicates that they are highly similar in the embedding produced by the unsupervised pipeline, which captures the most prominent signals defined by the top principal components. We argue that in order to separate these islet endocrine subtypes, more emphasis should be placed on the five hormone genes when performing cell embedding. To do that, we attached the GOI module to the bottleneck layer of SAKURA (Figure 2b), which takes the five hormone genes as knowledge input and encourages the cell embedding to separate cells with different expression levels of these genes (Figure 2b, Methods). When we used this knowledge-guided embedding of the extracted cells to run the same clustering algorithm used by the unsupervised pipeline, the resulting clusters displayed much more distinct expression patterns of the five hormone genes (Figure 2c). By further forming a hierarchy of these clusters based on their similarities, 11 cell groups are formed, all of which have unique expression patterns of the five hormone genes (Figure 2c).

In some of these 11 cell groups, multiple hormone genes are expressed. To make sure that this is not due to the presence of undetected sub-clusters, namely in a single cluster different subsets of cells express different hormone genes mutually exclusively, we examined individual cells in these clusters and found that most of them actually express multiple hormone genes (Figure S2). This is consistent with murine lineage tracing studies that showed that 1) virtually all hormone-expressing cells in the adult arise during fetal development and 2) multi-hormonal islet cells exist even in the adult ^28^. Because lineage tracing is not feasible in humans, analysis enabled by SAKURA can potentially be used to identify islet cell subpopulations throughout human development and determine how they are altered by disease, such as diabetes.

We further performed differential expression analysis between these 11 cell groups to identify genes most specifically expressed in one or a few of these groups (Figure S3). We found that these genes correlate with the expression of individual hormone genes but in a cluster-specific manner, rather than having a universal correlation with the expression of the hormone genes across all cells. For example, the expression of *PLAGL1*, *CCSER1*, *ACVR1C*, *PCSK1*, and *UNC5D* correlates with the expression of *INS*, but their co-expression is strongest only in 3 clusters that express only *INS* but not in other clusters that express *INS* together with other hormone genes. PCSK1 is a prohormone convertase required for proinsulin processing to insulin, while UNC5D regulates glucose-stimulated insulin secretion. PLAGL1 is a transcription factor that, when imprinted incorrectly, leads to beta cell loss and transient neonatal diabetes mellitus. Similarly, the expression of *KCTD8*, *CNTN5*, *NLGN1*, *LSAMP*, and *LEPR* correlates with the expression of *SST*, but their co-expression is strongest only in one cluster that expresses *SST* alone and one cluster that expresses only *SST* and *INS*. Interestingly, these 5 genes enriched in *SST* positive cells are mainly known for their roles in the nervous system. Overall, these results demonstrate SAKURA’s ability to characterize different groups of cells that fail to be separated in an unsupervised analysis, which can lead to an over-estimation of cells co-expressing multiple hormones (Figure 2a).

To more systematically evaluate the clusters produced from SAKURA’s embedding, we defined a distinctiveness measure to quantify how distinct the expression pattern of a gene is in different clusters (Methods). Using this measure, we found that clusters produced from SAKURA’s embedding display more distinctive expression patterns of the five hormone genes than clusters produced from cell embeddings of the standard unsupervised pipeline, four other unsupervised cell embedding methods, and a baseline version of SAKURA without the GOI module (Figures 2d, S4, S5). SAKURA’s results remained stable with different distance functions used in its loss formula as long as either L1 or L2 was employed in the GOI loss (Figure S5, Methods).

It is important to ensure that SAKURA’s knowledge-guided approach does not distort the embedding by focusing only on the five hormone genes while ignoring other genes. Data distortion in dimensionality reduction is often characterized by an artificial distribution of cell features in the embedding, which results in ambiguous clustering structures. Therefore, we evaluated cell clusters produced by the same clustering algorithm based on embeddings produced by SAKURA, the baseline version of SAKURA without the GOI module, and the five unsupervised methods. We assessed the quality of the clusters using three popular “internal validation” measures, which consider the compactness of each cluster and separation of different clusters without using any information about actual cell types. Based on these measures, SAKURA’s embedding ranks first or second among the seven methods in all cases (Figure S6a), showing that SAKURA’s attention to the five hormone genes did not lead to a strong distortion of the embedding. Again, SAKURA’s results were stable across different combinations of distance functions used in its loss formula (Figure S6b).

We also evaluated how SAKURA’s results would be affected by the presence and form of its regularization component and found that it consistently outperformed the standard unsupervised pipeline in all the settings tested (Figure S7). To make it easy to compare different methods, we developed a procedure to decide whether to include the regularization component when analyzing each data set based on the number of clusters produced (Methods).

### Knowledge-guided refinement of cell type annotations

Given that the embedding produced by SAKURA captures expression of the five hormone genes, we explored the use of it in two practical applications, namely determining cell types of previously unannotated cells and identifying cells whose previous annotations are suspicious.

For the first application, we took all cells from the pancreas data set and used SAKURA to produce an embedding, again with the five hormone genes supplied as knowledge input. In the corresponding UMAP plot, cells expressing the hormone genes are mainly concentrated in several areas (Figure 3a,b). In these areas, there are cells that the original authors of the cell atlas (Cao et al. ^27^) did not assign any cell type labels to (Figure 3a, “Unannotated”), and there are cells that were only given the broad label of “Islet endocrine cells” (Figure 3a). We extracted all cells in these areas, re-clustered them (Figure 3c), and set out to see if some of these cells can be annotated specifically as one of the islet endocrine subtypes.

**Fig. 3:**
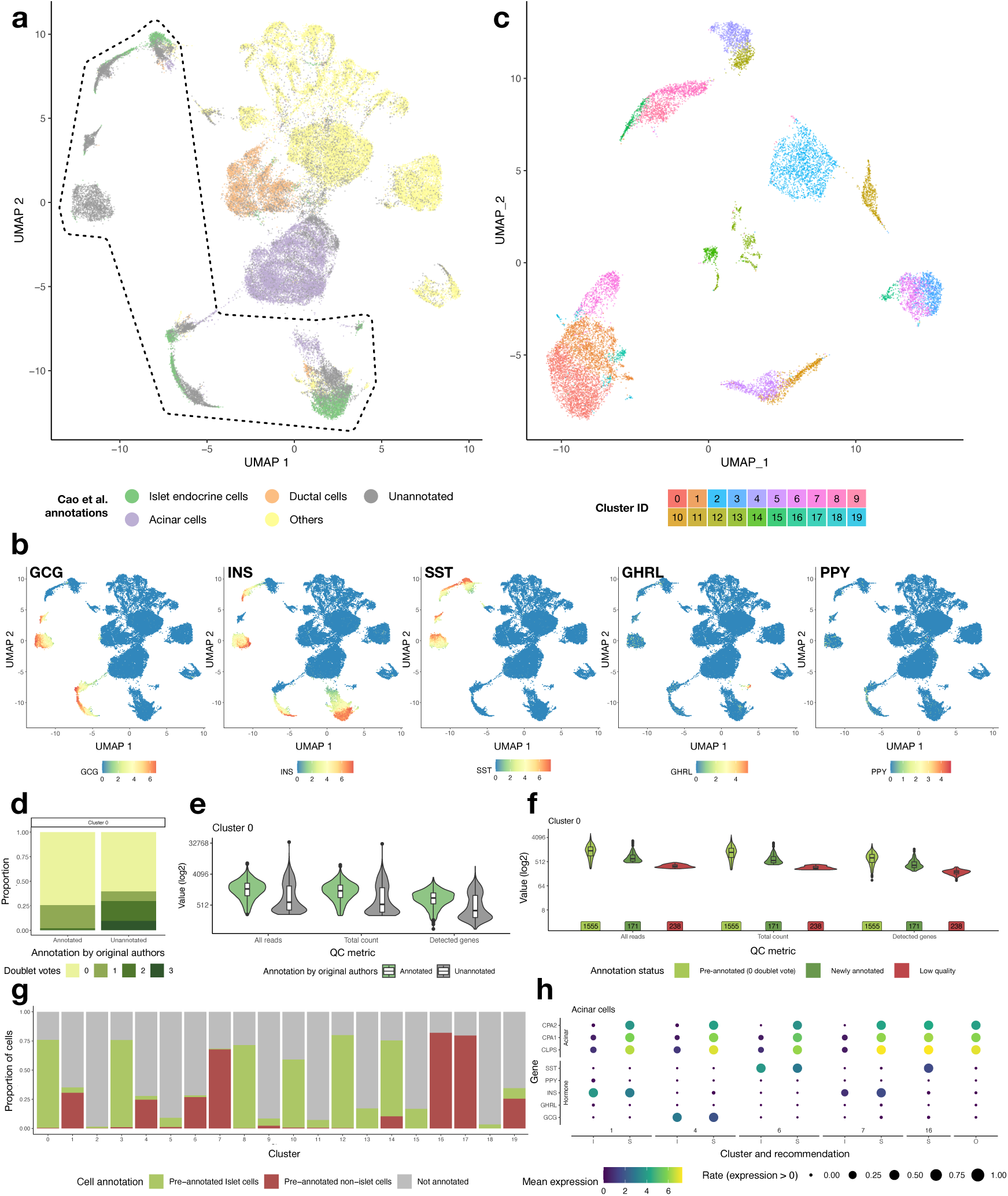
Applications of SAKURA in cell type annotations. **a,** UMAP visualization of all the cells in the pancreas data set based on SAKURA’s embedding. Cells are colored by their cell type labels according to the original authors’ annotations. The black contour line marks cells that express the five hormone genes, which form the “selected set”. **b,** Expression of the five hormone genes in different cells based on the same UMAP visualization as in Panel a. **c,** Re-clustering of the cells within the selected set based on SAKURA’s embedding. **d,** Number of doublet detection methods that vote for a cell as a doublet, for cells annotated or unannotated by the original authors. In Panels d, e, and f, data from cells in Cluster 0 are shown as examples. **e,** Quality measures of the cells annotated or unannotated by the original authors. **f,** Quality measures of the newly annotated cells as compared to other cells in the selected set. Cells in the “Pre-annotated (0 doublet vote)” group were annotated by the original author and not reported as doublets by any of the three doublet detection methods we used. Cells in the “Newly annotated” group are the originally unannotated cells in the selected set that passed our doublet and quality requirements. Cells in the “Low quality” group are the originally unannotated cells in the selected set that did not pass our doublet or quality requirements. **g,** Proportions of islet, nonislet, and unannotated cells in the hormone-expressing clusters based on the original authors’ cell type annotations. **h,** Expression of the acinar cell marker genes and hormone genes in the cells identified by SAKURA as requiring re-checking. I: cells annotated by the original authors as islet endocrine cells. S: cells annotated as acinar cells by the original authors that are marked as suspicious by SAKURA. O: cells annotated as acinar cells by the original authors that are outside the hormone-expressing clusters produced by SAKURA, which serve as positive controls of acinar cells.

We started by investigating the reason that some of these cells were not annotated by the original authors. We found that, on average, these unannotated cells have a higher chance of being doublets or having other data quality issues, but there are also some unannotated cells with low doublet probability and high data quality, as seen from the distributions of the QC metrics (Figures 3d,e, S8). Cells in this latter group were thus likely unannotated not because of quality issues but because they were not put into clusters with a clear expression of marker genes. We therefore developed a procedure to identify cells that i) were not annotated by the original authors, ii) have high data quality, and iii) are in one of the clusters identified by SAKURA that express only one of the hormone genes (Figure S9, Methods). We confirmed that these newly annotated cells have data quality resembling cells receiving cell type annotations from the original authors more than low-quality cells (Figures 3f, S8). Overall, our procedure identified 17 *α* cells, 171 *β* cells, 22 *δ* cells, and 38 *ε* cells among the cells that the original authors did not assign a cell type label to. Based on clusters that expressed only one hormone gene, we also gave specific endocrine subtype labels to 3,520 cells previously annotated by the original authors broadly as islet endocrine cells, including 379 *α* cells, 2,082 *β* cells, 629 *δ* cells, and 182 *ε* cells. Together, we assigned islet endocrine subtype labels to 3,520 cells in total (Supplementary File S1).

For the second application, we looked for cells in one of the islet endocrine cell clusters identified by SAKURA but were given a non-endocrine cell type label by the original authors. Some of these clusters have a high proportion of cells annotated by the original authors as other cell types (Figures 3g, highest in Clusters 1, 4, 6, 7, 16, and 17). We identified those with high expression of the five hormone genes and marked them as having suspicious cell type annotations (Figure S9, Methods, Supplementary File S2). For example, in the case of acinar cells (Figure 3h), some cells in these clusters annotated by the original authors as acinar cells (“S”) express the hormone genes at a level comparable to those annotated by them as islet endocrine cells (“I”) and much higher than cells annotated by the original authors as acinar cells that are outside these clusters (“O”). Since acinar cells are not typically known to express hormone genes at such levels, this suggests a potential ambiguity in their identity. This rationale also extends to other cell types, such as ductal cells, erythroblasts, and stromal cells (Figure S10). In total, we detected 1,833 cells whose original cell type annotations are suspicious. We emphasize that our intention in marking these cells as suspicious is not to assert misannotation by the original authors with certainty but rather to recommend verification of these annotations.

### Improved identification of CD4^+^ and CD8^+^ T cells

The pancreas data set analyzed above does not contain any cells annotated by the original authors with the five islet endocrine cell subtypes, and therefore we could not run cell embedding methods that require example cells of these subtypes as external knowledge input and compare SAKURA with them. In order to perform this type of comparisons, we next analyzed a single-cell data set produced from blood samples of COVID-19 patients ^29^ using CITE-seq ^30^. CITE-seq measures both transcript levels and cell surface protein levels using antibody-derived tags (ADTs). The original authors of this study combined transcript and ADT information to define cell types. We used these cell type annotations as ground truth to supply example cells to MCML and scNym, two semi-supervised methods that require example cells as external knowledge input. We then compared their cell embeddings with the embeddings produced by SAKURA, which only took gene names as knowledge input through its GOI module. All methods were given only the transcript levels of the data set but not the ADT information when producing their embeddings.

When we processed the data using a standard unsupervised analysis pipeline (Methods), CD4^+^T cells and CD8^+^ T cells, as defined by the original authors’ annotations based on both ADT and transcript information, largely mixed in the embedding (Figure 4a). In previous publications that involve analysis of single-cell transcriptomic data of immune cells, it is also common to find CD4^+^ and CD8^+^ T cells not clearly separated in the embedding (e.g., Ref ^31–33^).

**Fig. 4:**
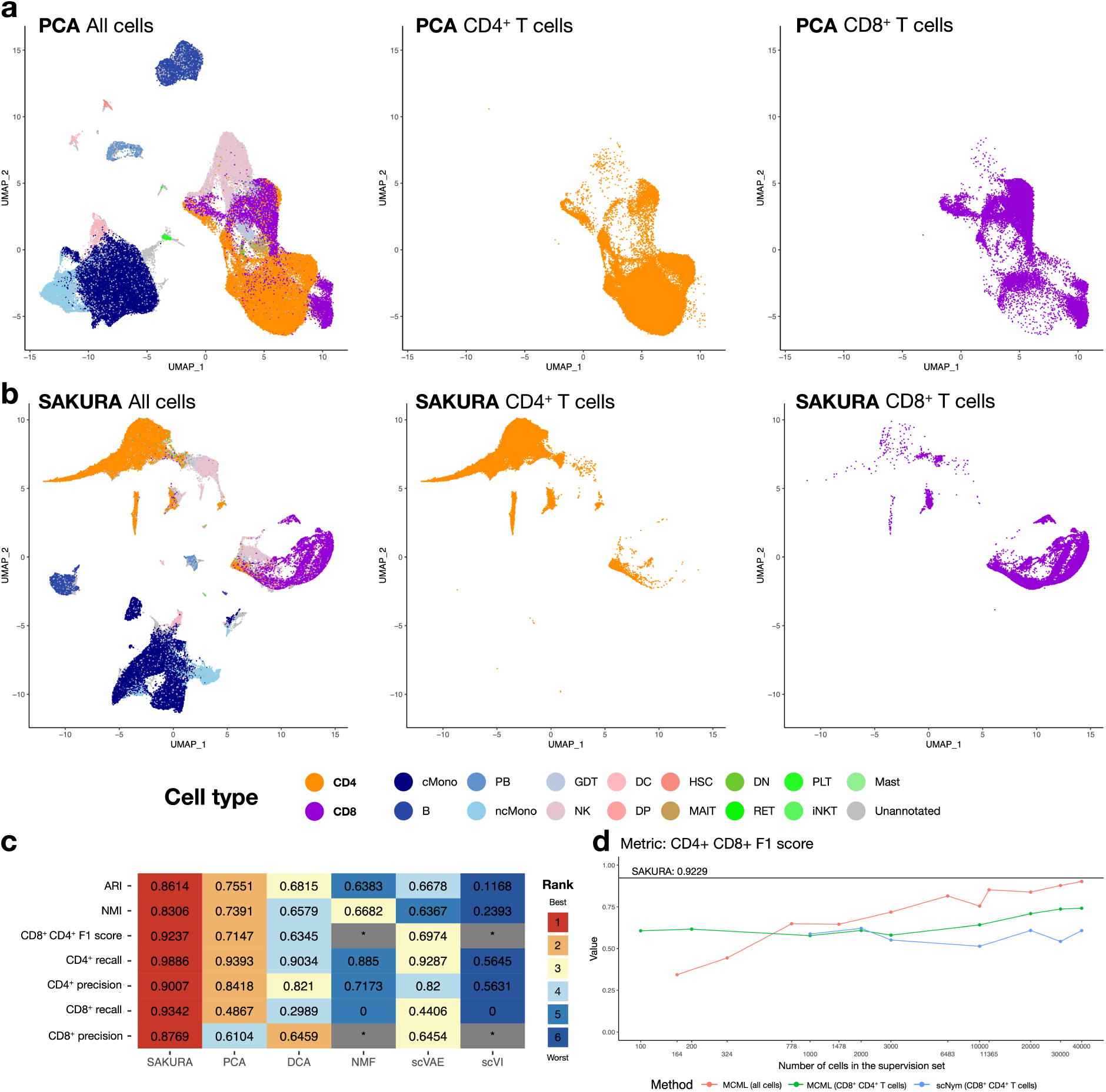
Improving separation of cell subtypes by SAKURA. **a,b,** UMAP visualization of the COVID-19 data set (all cells, only CD4^+^ T cells, or only CD8^+^ T cells) based on embedding produced by the unsupervised pipeline (a) or SAKURA (b) when they were supplied with transcript data only. **c,** Performance of the full-data set embeddings produced by SAKURA and five unsupervised methods in separating CD4^+^ and CD8^+^ cells with global cell type fidelity. Each row shows the actual performance scores and corresponding ranks of the methods based on one external evaluation measure. In cases that a measure could not be computed due to the clusters formed, the entries are marked by an asterisk. “CD4^+^ CD8^+^ F1 score” is the mean of the CD4^+^ F1 score and the CD8^+^ F1 score. **d,** Performance (F1 score) of the testing-set embeddings produced by SAKURA and two semi-supervised methods that take example cells as external knowledge input in separating CD4^+^ and CD8^+^ cells. The horizontal black line shows SAKURA’s performance. The three colored lines show performance of the two methods supplied with different numbers of cells as knowledge input, for all cell types or only CD4^+^ and CD8^+^ cells. All cell type labels represent external reference CITE-seq-derived cell type annotations (COVID-19 data set; Methods).

To see whether CD4^+^ T cells and CD8^+^ T cells can be better separated by our knowledge-guided approach, we supplied *CD4* and *CD8A* + *CD8B* as external knowledge to SAKURA’s GOI module (Methods). In the resulting embedding, CD4^+^ T cells and CD8^+^ T cells are more separated, and the CD4^+^ T cells further break into multiple clusters (Figure S11a). This is because some CD4^+^ T cells have low expression of *CD4* at the transcript level (Figure S11b) but they have a high level of CD4 at the surface protein level (Figure S11c). To focus on the separation of CD4^+^ T cells and CD8^+^ T cells without being affected by subsets of CD4^+^ T cells that express different levels of *CD4*, we reran SAKURA using only *CD8A* as knowledge input. In the resulting embedding, CD4^+^ and CD8^+^ T cells are better separated into different clusters (Figure 4b) than in the unsupervised case (Figure 4a). The small number of cells not clustered correctly were due to intrinsic limitations of transcript-level data. Specifically, some CD4^+^ T cells do not express *CD4* more highly than CD8^+^ T cells at the transcript level, and conversely some CD8^+^ T cells do not express *CD8A* and *CD8B* more highly than CD4^+^ T cells at the transcript level (Figure S12).

To quantitatively evaluate how well SAKURA’s embedding separates CD4^+^ and CD8^+^ T cells, we produced cell clusters and compared them with the authors’ cell type annotations. We also applied the same clustering method based on the cell embeddings produced by five unsupervised methods. To ensure a fair comparison with unsupervised methods, cell embeddings were evaluated using all cells in the entire data set. Quantified by seven evaluation measures (the “external validation” measures, Methods), SAKURA’s embedding consistently provided better separation of cell types than the embeddings produced by the other five methods in terms of both global cell type fidelity and the critical CD4^+^ and CD8^+^ T cell resolution (Figure 4c). In addition, based on the three internal validation measures and two ways to compute cell-cell distances, we found that SAKURA did not create stronger distortion to the embedding than the five unsupervised methods (Figure S13). To see whether SAKURA’s superior performance was solely due to the input marker genes, we also produced an embedding using only *CD4* and *CD8A* + *CD8B* transcript levels and then performed cell clustering. The resulting clusters were much less able to separate CD4^+^ and CD8^+^ T cells than the clusters formed from SAKURA’s embedding (Figure S14). This result shows that SAKURA was able to combine information about both the genes in the knowledge input and other genes when forming the embedding.

Next, we compared SAKURA with the two semi-supervised embedding methods. Based on the authors’ cell type annotations, we sampled cells from each cell type and supplied them as knowledge inputs to these two methods. We tested different amounts of example cells, in terms of either absolute cell number or percentage of all annotated cells, either for all 17 cell types in the data set, or for CD4^+^ and CD8^+^ T cells only from the training set (Methods). Then, we evaluated the embeddings of cells in the left-out testing set. We found that when more example cells were supplied, these two methods generally achieved better separation of different cell types (Figures 4d, S15). However, even when a very large number of example cells were supplied (e.g., 30,000), these methods still could not achieve better separation of different cell types than SAKURA (Figures 4d, S15). These results demonstrate the advantage of SAKURA that it requires only gene names as external knowledge input.

We performed three additional tests to evaluate the robustness of SAKURA. First, we conducted two additional runs of SAKURA’s embeddings, each using a different subsampled set of 200k cells from the COVID-19 data set. Each subsample was generated using a unique random seed to ensure variability in the selected cells (Methods). Compared to the cell embedding from the standard unsupervised pipeline, all SAKURA embeddings consistently showed enhanced global cell type fidelity and improved separation of the critical CD4^+^ and CD8^+^ T cells according to external validation measures (Figure S16). These results indicate that SAKURA consistently achieved similar performance across all three subsampled data sets, reinforcing the reliability of our findings.

Second, we modified the genes in the external knowledge input by either omitting some marker genes, using alternative marker genes, or including some irrelevant genes. In total, we tested 8 additional lists of knowledge input (Methods). We evaluated SAKURA’s embedding in each case by comparing the cell clusters formed with the original authors’ cell type labels. The results show that SAKURA’s embedding consistently separated CD4^+^ and CD8^+^ T cells better than the five unsupervised cell embedding methods we compared with in most settings (Figure S17a). Among the 8 lists, providing only *CD4* as knowledge input led to slightly worse performance as expected due to some CD4^+^ T cells that do not express *CD4* at the transcript level, as explained above, as was the case when two other marker genes of CD4^+^ T cells were supplied as the only knowledge input, *CCR7* and *IL7R*. Importantly, even in these cases, SAKURA’s performance was still better than or comparable to the other five methods. Also, including genes not expected to be useful for separating CD4^+^ and CD8^+^ T cells (*IFNG*, *LYZ*, or *FN1*) did not result in a major performance drop.

We also used the internal validation measures to check whether SAKURA’s embedding was distorted by the knowledge inputs. As compared to the five unsupervised methods, SAKURA’s embedding was not more strongly distorted regardless of the genes on the list of knowledge input (Figure S17b).

In the third additional test, we compared different settings of the supervision intensity parameters *λ*_4_, *λ*_5_, *λ*_6_, which are used to specify the relative importance of the external knowledge input (Methods, Figure S18). In terms of separating CD4^+^ and CD8^+^ cells, SAKURA performed better than all the unsupervised methods for a wide range of intensity values around the default setting (Figure S18a, from “X0.0001_0.01” to “X1_100”) according to all three performance measures. At these supervision intensity settings, the embeddings produced by SAKURA were also not more distorted than the embeddings produced by the unsupervised methods (Figure S18b). Importantly, when the supervision intensity was set too strong, the AWS measure clearly signaled the data distortion problem by having negative values (Figure S18b, from “X10_1000” to “X1000_100000”), which provides a practical way to detect and avoid over-supervision when the external validation measures cannot be computed due to lack of externally acquired cell annotations. Together, these results show that SAKURA led to robust performance across a wide range of supervision intensity parameter values, and suitable values of these parameters could be determined based on the sign of the AWS measure.

### Identification of monocyte sub-population that correlates with COVID outcome

Previous studies have underscored the pivotal role of the S100A protein family in modulating inflammatory responses in different disease contexts, such as cancer and cardiovascular disease ^34,35^. Concurrently, using the ADT information of the COVID-19 data set, the original authors discovered a cell cluster that contains S100A8/9/12^hi^ HMGB2-expressing monocytes and found that patients with a larger proportion of cells coming from this cluster exhibited increased illness severity ^29^. We found that this proportion also correlates with disease outcome (Figure 5a), which is another important clinical indication. To determine whether similar findings could be obtained using transcript data alone, since ADT information is not available in standard single-cell transcriptomic data, we used a standard unsupervised data analysis pipeline to produce cell embedding and cluster cells based on only the transcript portion of the data set. Using this standard approach, we were only able to find a large cluster of monocytes that is roughly divided into a subpopulation of S100A8 and S100A9 expressing classical monocytes and a subpopulation of non-classical monocytes (Figure 5b,c). The proportion of classical monocytes only moderately correlates with COVID outcome (Figure 5d).

**Fig. 5:**
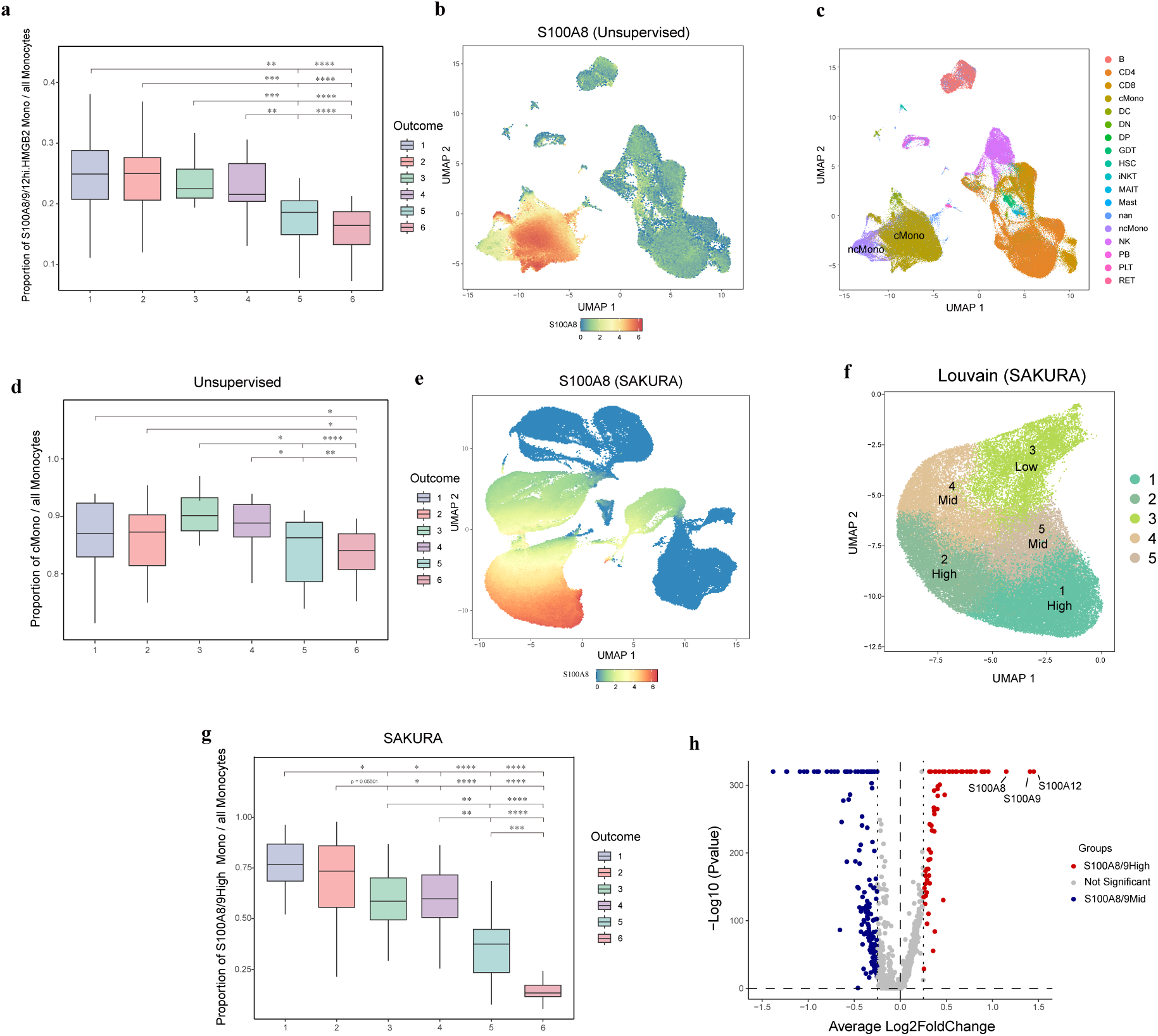
Identifying clinically relevant cells using SAKURA. **a,** Proportions of S100A8/9/12hi.HMGB2 monocytes, identified by combining transcript and ADT information by the original authors, in patients with different COVID outcomes (smaller values for more serious outcomes. 1: death; 2: intubated, ventilated; 3: non-invasive ventilation; 4: hospitalized, O2; 5: hospitalized, no O2; 6: not hospitalized). Each box plot summarizes the cell proportions in different patients with the same outcome. **b,c,** UMAP plot of cells based on the embedding produced by the unsupervised pipeline applied to transcript data only, with cells colored by the expression level of *S100A8* (b) or classical monocytes (cMono) and non-classical monocytes (ncMono) annotated by the original authors (c). **d,** Proportions of classical monocytes identified by the unsupervised pipeline in patients with different COVID outcomes. **e,** UMAP plot of cells based on the embedding produced by SAKURA applied to transcript data only, with cells colored by the expression level of *S100A8*. **f,** Lovain clusters of monocytes identified from the embedding produced by SAKURA, which express *S100A8* and *S100A9* at high (High), medium (Mid), or low (Low) levels. **g,** Proportions of S100A8/9hi monocytes identified by SAKURA in patients with different COVID outcome. **h,** Differentially expressed genes between the two classical monocyte subpopulations, S100A8/9High and S100A8/9Mid. In Panels a, d, and g, statistical significance was determined by a one-sided Wilcoxon rank-sum test of the proportions in two groups. *: p*<*0.05; **: p*<*0.01; ***: p*<*0.001; ****: p*<*0.0001.

We suspected that the low correlation was due to imperfect separation of monocytes with different expression levels of S100A8 and S100A9. Therefore, we used SAKURA to produce cell embedding with both S100A8 and S100A9 supplied as external knowledge input through its GOI module and compared that with the embedding from the unsupervised method (Figure S19). In the embedding produced by SAKURA, there was a cluster of monocytes with a clear gradient of expression of S100A8 and S100A9 from top to bottom of the cluster (Figure 5e). This allowed us to form different monocyte sub-clusters that express different levels of these genes (Figure 5f). The proportion of monocytes that belong to the S100A8/9hi cluster correlates very well with COVID outcome (Figure 5g), showing that when ADT information is not available, clinically important cell clusters can be recovered from transcript levels only by providing key marker genes as knowledge input to guide the embedding process.

We further performed a differential expression analysis between the classical monocytes in S100A8/9hi and S100A8/9mid clusters. We found a list of genes significantly differentially expressed between the two groups (Figure 5h), showing that cells in the two groups are consistently different in the expression of these genes, thus supporting that the two groups of monocytes were not artificially created by SAKURA based on expression levels of *S100A8* and *S100A9* solely due to our knowledge input.

### Identification of rare senescent cells

As the third demonstration of SAKURA, we used it to identify rare senescent cells using a single-nucleus RNA-seq (snRNA-seq) data set of young and old mouse brains ^36^. The characteristics and roles of senescent microglia in the aging brain are under active investigation ^37^. To see if these cells can be identified from the snRNA-seq data set, we extracted all the microglia and performed a standard unsupervised analysis, which produced a number of cell clusters (Figure 6a). When we checked the expression of the cyclin-dependent kinase inhibitors (CDKI), which is a hallmark of cellular senescence ^38^, we did not find microglia that highly express CDKI to be clustered (Figure 6b). Similarly, when we examined the expression of genes indicative of molecular aging ^39^, microglia with high expression of these genes were scattered (Figure 6c). These results suggest that information about senescent cells was not among the major defining features of the embedding produced by the unsupervised pipeline.

**Fig. 6:**
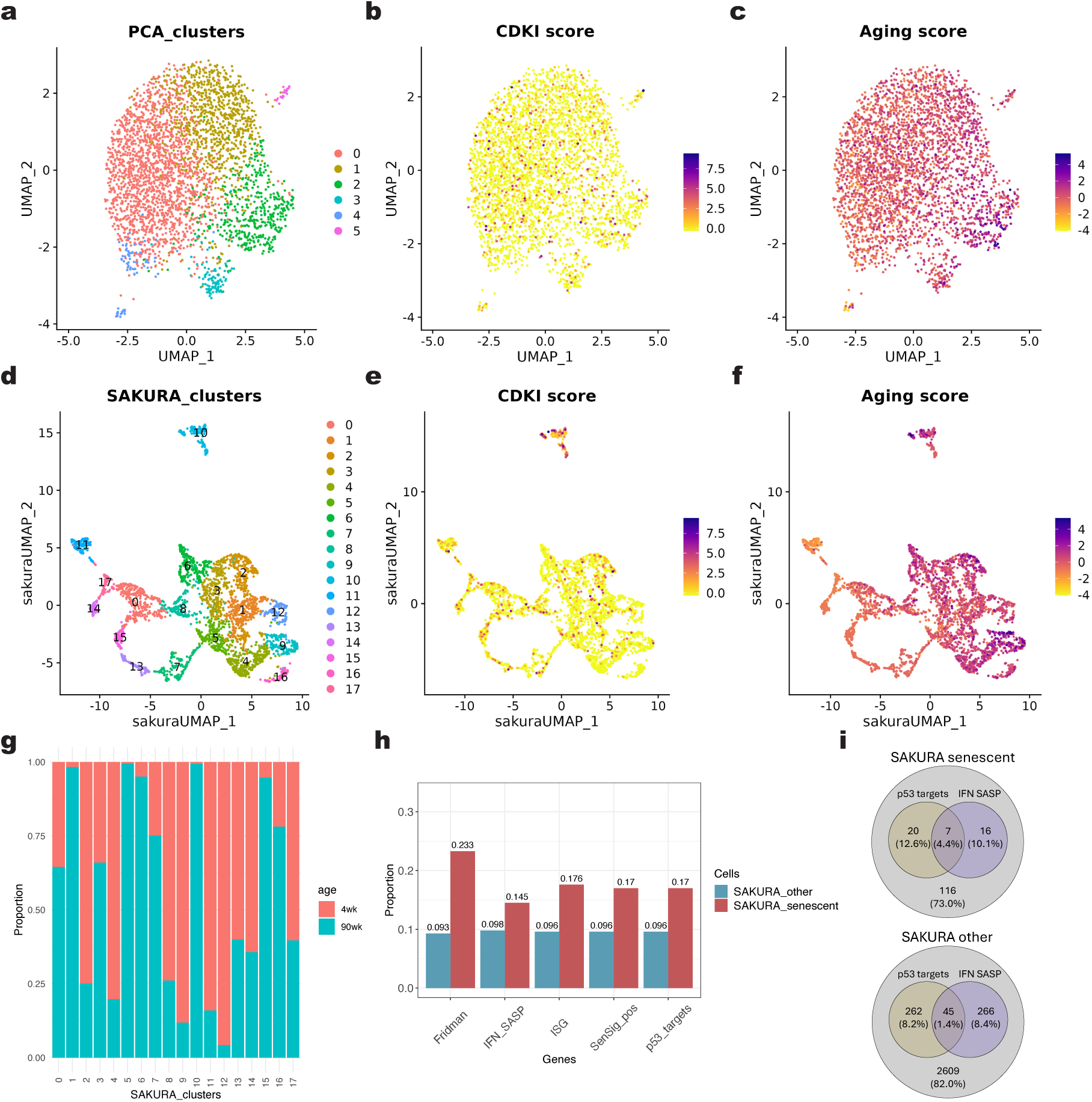
Identifying senescent cells using SAKURA. **a,** UMAP visualization of mouse microglia cells in the aging data set based on the unsupervised embedding. Cells are colored by clustering results using the unsupervised pipeline. **b,c,** UMAP plots showing CDK inhibitor scores (b) and aging scores (c) based on the unsupervised embedding. **d,** UMAP visualization based on SAKURA’s embedding. Cells are colored by clustering results using SAKURA’s embedding. **e,f,** UMAP plots showing CDK inhibitor scores (e) and aging scores (f) based on the SAKURA’s embedding. **g,** Proportion of cells from young (4 weeks) and old (90 weeks) samples in Cluster 10. **h,** Proportion of cells that have top 10% scores calculated using genes from five categories (genes up-regulated in senescent cells proposed by Fridman and Tainsky ^69^, genes associated with either interferons (IFNs) or senescence-associated secretory phenotype (SASP) [manually curated], interferon-stimulated genes (ISGs) ^70^, genes with positive coefficients in the SenSig gene set ^71^ and p53 target genes ^72^), in the potential senescent cells identified using SAKURA (Cluster 10, “SAKURA_senescent”) and the other cells (“SAKURA_other”). **i.** Venn diagrams showing the number and percentage of cells with top 10% scores calculated using IFN-SASP genes and p53 target genes, in both potential senescent cells (top) and all other cells (bottom) identified using SAKURA.

Next, we ran SAKURA with the CDKI and molecular aging signature genes supplied as knowledge inputs (Methods). In the resulting cell clusters (Figure 6d), Cluster 10, which constitutes 4.76% of all microglia, is enriched in cells expressing both CDKI and molecular aging genes strongly (Figures 6e,f, S20). Consistently, the majority of cells in this cluster came from old mice (Figure 6g). As further support that microglia in Cluster 10 are senescent cells, we found a larger proportion of them highly expressing several other senescence-related gene sets than cells in other clusters (Figure 6h). In addition, cells in Cluster 10 also co-expressed multiple senescence signatures more frequently than the other cells (Figure 6i). These results demonstrate that by supplying partial knowledge of some cells (CDKI and aging signatures), the embedding produced by SAKURA can facilitate identification of cells of interest and studying of their other properties (senescence-associated secretory phenotype [SASP], p53 targets, etc.).

## Discussion

Despite these encouraging empirical results presented, there may still be a concern that SAKURA is not an “unbiased” approach. Our response to this putative concern is three-fold.

First, we emphasize that SAKURA does not replace standard unsupervised methods but instead provides an alternative when these methods are not suitable. When the goal of analysis is to retain the most prominent signals in the data, such as major cell types, indeed an unsupervised method can work well. However, if some biologically important signals cannot be captured by an unsupervised method, the genes of interest taken by SAKURA as external knowledge input can help define the analysis goal by modifying the loss function used by the model training process.

Second, it should be noted that in machine learning, inductive bias is always necessary to make learning possible ^40^. Inductive bias is the set of assumptions made to enable predictions for unseen instances, which manifests in the representation of the possible models and the corresponding learning algorithm. Therefore, even an unsupervised method incurs “biases”, such as the assumption that cells of a cell type cluster together and are separated from other cells in the space formed by the most variable genes. Sometimes these biases are not explicitly stated and sometimes the user is not aware of them, but they do actually exist.

Third, when using some unsupervised dimensionality reduction methods, the optimization problem involved can be one that does not have a unique solution or has different parameterizations that give similar loss values. The external knowledge input of our knowledge-guided approach can help overcome these problematic situations.

Our analysis of the fetal pancreas data suggests that SAKURA can be used to address important questions in islet biology requiring extremely precise determination of cell sub-populations. Some of these questions include 1) whether beta-cell maturation from human pluripotent stem cells mimics the normal fetal developmental program and 2) whether the dedifferentiation or transdifferentiation of islet cells from patients with diabetes reflects reversion to a state observed in fetal development.

Although in this study we have focused on one single type of knowledge input, namely a set of genes of interest, the modular design of SAKURA also allows it to employ other types of knowledge and their combinations. For example, if example cells of a rare cell type are available, they can be supplied in combination with a set of marker genes of this cell type as two separate types of knowledge input. Supplying multiple types of knowledge can help overcome the limitations of each individual type.

## Conclusions

In this study, we have demonstrated that the knowledge-guided approach of SAKURA can help distinguish among highly similar cell types (e.g., islet endocrine cells and hematopoietic subpopulations) and identify rare cells (e.g., senescent cells). As compared to existing knowledge-guided cell embedding methods for single-cell transcriptomic data, which all require labeled example cells as knowledge input, SAKURA only requires the names of some genes of interest as external knowledge, which are much easier for the user to provide. In addition, we have also demonstrated that, as reflected by several internal validation measures, the knowledge-guided approach of SAKURA does not lead to stronger distortion of the embedding space as compared to unsupervised methods that do not take external knowledge as input. Moreover, we have demonstrated that even if the input genes of interest are noisy and do not contain complete knowledge, they can still effectively guide the cell embedding process to aid discovery of important rare signals.

## Methods

### Details of SAKURA

SAKURA is a flexible framework that allows the attachment of different modules for taking different types of knowledge input to guide the cell embedding process. Each module is in the form of an additional functional unit added to the autoencoder backbone, and correspondingly an additional component added to the loss function to reflect which embedding is not preferred according to the external knowledge supplied. The relative importance of the knowledge input as compared to the overall data properties is controlled by supervision intensity parameters.

#### The autoencoder backbone

Denote the processed data as D, which is an *M* × *N* matrix of log-transformed UMI counts of *N* features (i.e., genes) in *M* cells. For the gene expression profile *x_i_* of any cell *i*, the encoder *f^enc^*(*x_i_*) = *h_i_* generates a low-dimensional representation *h_i_* of it, while the decoder *f^dec^*(*h_i_*) = *x*^′^ attempts to regenerate *x_i_* from *h_i_*. From the perspective of preserving useful information that can reconstruct the input, a good autoencoder is one that has the output of the decoder, *x*^′^, similar to the input, *x_i_*. We quantified this by defining a reconstruction loss *L^rec^* that considers L1 and L2 distances ^41^:

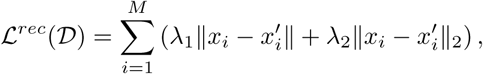

where *λ*_1_ and *λ*_2_ are hyperparameters that control the weights of the two distance terms.

In our current implementation, both the encoder and decoder are composed of three hidden layers of nodes, each in the form of a linear transformation followed by the CELU activation function. For the pancreas use case, we used 100 nodes per layer, while for the COVID-19 and aging use cases, we used 200 nodes per layer due to the larger data size. The bottleneck layer between the encoder and the decoder always has 50 nodes.

#### Regularization

To increase numeric stability during the training process, we add a regularization component, *L_reg_*, to the loss function:

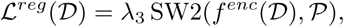

where SW2(·) is the sliced 2-Wasserstein distance ^42^, P is a predefined distribution, and *λ*_3_ is a hyperparameter. SW2(·) projects both the embedding distribution and the predefined distribution onto multiple random one-dimensional spaces (called the slices) and then uses Wasserstein distance to quantify their discrepancy. We chose to use SW2 because by capturing distribution geometry through optimal transport ^43^, it maintains structural relationships even when there is a distribution mismatch ^42^. In addition, with a time complexity of O(*d* · *n* log *n*), where *d* is the number of dimensions in the embedding (set to 50 by default), SW2 is scalable to a large number of embedding dimensions. In comparison, the potential alternatives KL divergence and total variance could suffer from sensitivity to initialization or poor scalability ^44,45^.

By default, we set P ∼ Uniform(−*k, k*), which is a uniform distribution in all dimensions with *k* being an arbitrary constant. The uniform distribution is chosen to prevent representation collapse and encourage efficient utilization of the latent space. While the task-specific loss naturally clusters cells with similar expression data, the uniform constraint prevents the entire embedding from collapsing to a small region. This balance results in embeddings where distinct cell groups form separated clusters while the overall distribution occupies a broad and meaningful volume in the latent space.

#### Procedure for deciding whether to include the regularization component

As explained above, conceptually including the regularization loss can prevent the cells from being highly condensed in the embedding. In practice, we have found that sometimes even without regularization, SAKURA could still produce reasonable embeddings. In that situation, we would prefer to decide whether to include the regularization component based on the resulting number of cell clusters produced. Specifically, in our procedure, the regularization component would be included if and only if the number of clusters produced was closer to that of the standard unsupervised method (PCA embedding with 50 dimensions, followed by Louvain clustering with default resolution). This procedure was applied to the analysis of all three data sets in this study.

#### Module for taking genes of interest as external knowledge input

The module that takes genes of interest (GOI) as knowledge input is a regression unit attached to the bottleneck layer of SAKURA’s backbone. It uses each cell’s embedding to infer expression levels of the GOI in the cell. Performance of the regression is evaluated by combining L1, L2, and cosine distances, which together form a component of the loss function, *L^goi^*:

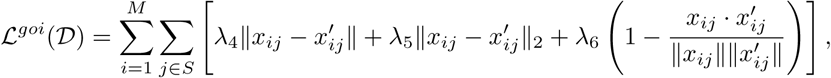

where *S* is the set of genes of interest, *x_ij_* is the expression level of gene *j* in cell *i*, *x*^′^ = *f^goi^*(*f^enc^*(*x_i_*)) is the inferred expression level of gene *j* in cell *i*, and *λ*_4_, *λ*_5_, and *λ*_6_ are hyperparameters. For the regression function *f^goi^*, we used a three-layer fully connected network with 50 nodes in the two hidden layers and one node per GOI in the output layer (Figure 2b).

This design can be easily generalized to handle arbitrary functions of gene expression levels. For example, in our application of SAKURA to separate CD4^+^ and CD8^+^ cells, we used the sum of *CD8A* and *CD8B* expression levels as the regression target since these genes encode the two chains of the CD8 receptor.

#### Overall loss function

Combining the three components, the overall loss function of SAKURA is as follows:

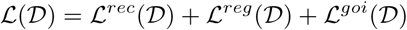

#### Model training and hyperparameter settings

To train the SAKURA model, we first randomly split the whole data set into a training set (50% of all cells) and a testing set (remaining 50% of the cells). We used the batched gradient descent method of the RMSProp optimizer ^46^ (with parameters *lr* = 0.001, *α* = 0.9) to perform model training based on the training set. In each iteration, we first optimized for *L^rec^* and *L^reg^* and then optimized for *L^goi^*. When optimizing for *L^rec^* and *L^reg^*, we used a batch size of 200 cells randomly sampled from the training set. The hyperparameters *λ*_1_ and *λ*_2_ were fixed at 1 during the whole training process, while *λ*_3_ was initialized to 0.0001 and got ramp-up by 1% per epoch with a cap at 1. This design allows a less constrained exploration of the parameter space at the beginning and stronger regularization at the end. When optimizing for *L^goi^*, we used a batch size of 100 cells randomly sampled from the training set. The hyperparameters *λ*_4_, *λ*_5_, and *λ*_6_ were either fixed to 0, or initialized to 0 and got ramp-up by 0.01 starting from the 50th epoch with a cap at 1, depending on whether each of the loss terms was included in the specific configuration tested. This setting was to let the model capture general properties of the data set first and progressively pay more attention to the genes of interest.

We use the testing set as a canary to detect any abnormal behaviors during the training process or data distortion in the resulting embeddings in two ways. First, we monitored the loss curves based on training data and testing data. Second, we compared the distribution of cells in the training and testing sets in the embedding. Having substantial difference in either case would indicate that the training process is problematic. In all the results presented in this manuscript, we did not detect any problems of the training process in this way.

#### Software availability

SAKURA is freely available as an open-source tool at https://github.com/Yip-Lab/SAKURA.

### Data collection and processing

#### Pancreas data set

For the Human Fetal Cell Atlas pancreas single-cell RNA sequencing data set, we obtained the raw UMI count matrix and cell type annotations provided by the original authors ^27^ from https://descartes. brotmanbaty.org/bbi/human-gene-expression-during-development/. We used Seurat (version 3.2.0) ^47^ to perform minimal pre-processing of the data, including log-transformation of the raw UMI counts using NormalizeData(), selecting the 10,000 genes with highest variance using FindVariableFeatures(), and scaling the data based on the high variance genes using ScaleData().

We then applied a standard unsupervised analysis pipeline to the data. It first performed dimensionality reduction using an unsupervised method, which was principal component analysis (PCA) by default. The embedding was then used as input to generate two-dimensional (2D) UMAP visualizations using RunUMAP(), using Euclidean distance as the distance metric by default. These PCs were used to generate a shared nearest neighbor (SNN) graph based on cell-cell Euclidean distance using Find-Neighbours(), which was then used to perform Louvain clustering using FindClusters() with the default resolution.

#### COVID-19 data set

For the COVID-19 CITE-seq data set, we obtained raw UMI counts and cell surface protein readings produced by the COVID-19 Multi-Omic Blood ATlas (COMBAT) consortium ^29^ from https://zenodo.org/records/6120249. We used the data in the “COMBAT-CITESeq-DATA” archive in this study. Since the total number of cells in this data set is large (847k) and some cell embedding methods involved in our comparisons could not handle it, we randomly subsampled 200k cells for all our analyses of this data set. Next, we used a standard procedure to process the data. Specifically, we extracted the raw count matrix of the transcriptome data and ADT features (“X” object) and the annotation data frame (“obs” object) from the H5AD file. We dropped all ADT features (features with names starting with “AB-”) and put the transcriptome data along with the annotation data frame into Seurat (version 4.1.1).

We then applied the same data processing and standard unsupervised analysis pipeline as the one we applied to the pancreas data set. Finally, we extracted the clustering labels generated and concatenated them with cell type, major subtype, and minor subtype annotations provided by the original authors, which were manually curated using both surface protein and transcriptome information.

To enhance the robustness of our analysis, we performed repeated processing with three distinct sets of subsampled 200k cells from the COVID-19 data set. The subsample random seed used for each run is as follows:

- Run 1 (main result): random seed = 114514
- Run 2: random seed = 1919
- Run 3: random seed = 810

By documenting these seeds, we clarify the random selection process and thereby facilitate future replication of our study.

#### Aging data set

For the aging snRNA-seq data set of young and old mouse brains ^36^, we obtained the processed data from https://cellxgene.cziscience.com/collections/31937775-0602-4e52-a799-b6acdd2bac2e. We selected cells annotated as microglia by the original authors. We applied the same data processing and standard unsupervised analysis pipeline used for the pancreas data set. The only modification made was selecting the 5,000 genes with highest variance using FindVariableFeatures(), since we focused on microglia only.

To identify senescent cells using SAKURA, we used a combination of cyclin-dependent kinase (CDK) inhibitor genes and aging genes. Specifically, we calculated CDK inhibitor scores by first averaging the z-scores of the log-normalized expression levels of seven CDK inhibitor genes (*Cdkn1a*, *Cdkn1b*, *Cdkn1c*, *Cdkn2a*, *Cdkn2b*, *Cdkn2c*, and *Cdkn2d*) and then calculating the z-score of this average. Similarly, we calculated aging scores using 82 aging signature genes identified in mouse brains as reported by Hahn et al. ^39^. To identify cells with both high CDK inhibitor scores and aging scores, we used the minimum of these two scores as the regression target of SAKURA.

### Other cell embedding methods compared

We compared SAKURA with different cell embedding methods, including unsupervised methods, which do not take external knowledge inputs, and semi-supervised methods, which take example cells as knowledge inputs.

#### Unsupervised cell embedding methods

We selected five single-cell embedding methods, including a standard method for dimensionality reduction (PCA), a matrix factorization method (NMF), and three methods based on autoencoder (DCA, scVAE, and scVI), which is a current mainstream approach. When applying the unsupervised methods, for each data set, we used data from all cells to generate cell embeddings.

1. Principal Component Analysis (PCA). PCA ^48,49^ is a classical dimensionality reduction method that finds the most variable directions in the data called the principal components (PCs), which are linear combinations of the original variables, with all PCs orthogonal to each other. We used the R package of Seurat (v3.2.0) ^47^ to perform PCA on the pre-processed gene expression data using its RunPCA() function. By default, we used the top 50 PCs to form the 50-dimensional cell embedding for downstream analyses.
2. Non-negative matrix factorization (NMF). NMF ^50^ is a method that decomposes a data matrix into the product of two matrices with no negative elements, respectively called the basis matrix and the encoding matrix, with a goal to maximally reconstruct the original data matrix. Dimensionality reduction is achieved by having fewer dimensions in the basis matrix than the original data. The non-negativity requirement makes interpretation easier. For example, for gene expression data, the original transcriptome profile may be explained by the superposition of a number of gene expression programs. We used the Python package sklearn (v1.3.1) ^51^ to perform NMF on the pre-processed gene expression data with 50 components. We used the 50 output features to form the 50-dimensional cell embedding for downstream analyses.
3. Single-cell variational autoencoder (scVAE). scVAE ^9^ is a method that uses variational autoencoders to estimate the expected gene expression levels based on the observed raw counts, and produce a latent representation for each cell during the process. We used the Python package scvae (v2.0.0) to produce cell embeddings with the following settings: GMVAE model with a negative binomial likelihood function, a 50-dimensional latent variable as cell embedding, two hidden layers of each of the 200 units, and 200 epochs using the warm-up scheme is trained for 500 epochs on the raw count data.
4. Deep Count Autoencoder (DCA). DCA ^8^ is a method that uses an autoencoder to denoise single-cell transcriptomic data, taking into account the overdispersion and sparsity of the count data, and nonlinear gene-gene dependencies. We used the Python package dca (v0.3.1) to train a deep count autoencoder network and produce a hidden representation of the cells. In the network, we used three hidden layers with 100, 50, and 100 neurons, respectively. We trained the network for 300 epochs to learn the 50-dimensional cell embeddings for downstream analyses.
5. Single-cell Variational Inference (scVI). scVI ^6^ is a method that uses stochastic optimization and deep neural network to find similar cells and aggregate their information in order to estimate the distribution of expression values. Batch effects and limited detection sensitivity are also taken into account. We used the Python package scvi-tools (v1.0.4) ^52^ to perform dimensionality reduction on the raw count data with model scVI. We extracted the 50-dimensional latent space representation as the cell embedding for downstream analyses.

#### Semi-supervised cell embedding methods

For semi-supervised methods, we used the same training and testing sets defined for training the SAKURA model, with all example cells required by these methods as input chosen from the training set. We tested the performance of the different methods when they were supplied with different amount of example cells in terms of absolute number (50, 100, 500, or 1000 cells per cell type) or percentage (1.5%, 5%, 10%, 15%, or 20% of cells of each cell type). When we supplied an absolute number of example cells as knowledge input, for cell types with fewer cells than required, all cells from these cell types in the training set were provided as input to the semi-supervised methods.

We selected two semi-supervised methods for our comparisons, MCML and scNym, which were the only semi-supervised methods developed for single-cell transcriptomic data that we were able to run. We also tested netAE ^18^ but it consumed prohibitive amounts of memory when running on the data sets we tested and thus we were not able to include it in our comparisons.

1. Multi-Class Multi-Label (MCML). MCML ^17^ is a method that uses an autoencoder to produce a cell embedding, with the cell type labels of the example cells involved in defining the loss function. We used the Python package MCML (v0.0.1) to produce cell embeddings. Following the default settings, we first centered the pre-processed gene expression data before using MCML to perform dimensionality reduction. Cell type annotations were then inferred using k-Nearest Neighbors (KNN) classification with K=30.
2. scNym. scNym ^19^ is a method that uses adversarial neural networks to transfer cell type annotations from one experiment to another. During the process, a cell embedding is learned by considering both the unlabeled cells and the cell type labels of the example cells. We used the Python package scnym (v0.1.10) to perform cell embedding. We adapted the “new_identity_discovery” configurations to handle the cell types that were excluded or unseen in the testing set, and trained the scNym model with two 50-unit hidden layers to extract the activations of the penultimate neural network layer as cell embeddings.

### Evaluation measures

We used a variety of measures to evaluate cell embeddings produced by SAKURA and other methods.

#### The distinctiveness measure for evaluating distinct gene expression in cell clusters

Dot plots are commonly used to visualize gene expression in cell clusters ^53^. In a dot plot, the size of a dot indicates the detection rate of a gene in a cluster (i.e., the proportion of cells with a non-zero expression of the gene) and the color of a dot indicates the average expression level among cells that express the gene. We used these two types of information to define a measure that quantifies how distinct the expression of a gene is in a cluster. Basically, it is desirable to have a gene distinctively expressed in one or a small number of clusters, and the degree of distinct expression depends on both the detection rates and the average expression levels of the gene in different cell clusters. Specifically, we define the distinctiveness score as follows. For gene *i* that has a detection rate of *t_ij_* in cluster *j*, defined as the proportion of cells in cluster *j* with a non-zero detected expression level of gene *i*, we define the detection distinctiveness score as

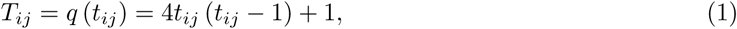

where *q*() is a quadratic function that gives a high score to a detection rate close to 1 and 0, while a low score to a detection rate close to 0.5. *T_ij_* has a range of 0 to 1 and a higher score indicates a situation closer to having all cells in cluster *j* expressing gene *i* or all cells in cluster *j* not expressing gene *i*. When this score is averaged over all clusters, a higher score indicates a situation closer to having gene *i* distinctively expressed in certain clusters and not expressed in other clusters.

We also create a similar score for the average expression level. Instead of using the absolute level, which varies from gene to gene, we use its empirical cumulative distribution function (ECDF) based on the expression level of a gene in all cells. Specifically, for gene *i* that has an average expression level of *e_ij_*in cluster *j*, we define the expression level distinctiveness score as

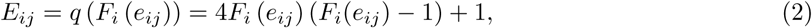

where *F_i_*() is the ECDF of gene *i*. *E_ij_* has a range of 0 to 1 and a higher score indicates a situation closer to having all cells in cluster *j* expressing gene *i* highly or having all cells in cluster *j* expressing gene *i* lowly, as indicated by an average expression level *e_ij_* closer to the 0th-percentile or the 100th-percentile according to the ECDF, respectively.

Based on the detection distinctiveness score *T_ij_* and the expression level distinctiveness score *E_ij_*, we define an overall distinctiveness score *G* by multiplying the two scores, averaging over all clusters, and further averaging over all genes of interest (e.g., the 5 hormone genes in the pancreas study or one of them) if a single score is needed:

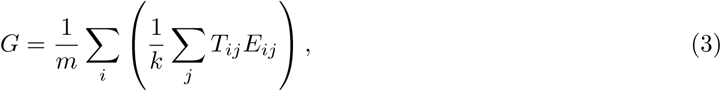

where *m* is the total number of genes of interest and *k* is the total number of cell clusters. The distinctiveness measure *G* has a range of 0 to 1, and a larger value indicates more distinct detection and expression levels of the genes of interest in the cell clusters.

A major advantage of the distinctiveness score over standard differential expression analysis is that it is much less affected by the number of clusters. For example, if a cell type is split into multiple cell clusters by a clustering algorithm and cells in all these clusters express a marker gene, in a standard differential expression analysis, the gene may not be significantly more highly expressed in any one cluster than in all the other clusters combined. In contrast, the distinctiveness score of the gene can still be high in each of these clusters. In principle, the distinctiveness score can be applied no matter how cells are grouped, such as by cell clusters through a clustering algorithm or by cell types based on a specific annotation procedure.

#### Internal validation measures for evaluating compactness and separation of cell clusters

We used three popular measures to evaluate cell clusters based on the compactness of each cluster and the separation of different clusters. These measures do not rely on an externally-acquired reference that defines how the cells should be clustered and therefore they are called “internal validation measures”. All three measures require a distance function *d*() in the embedding. We used both Euclidean distance and cosine distance in our comparisons, although some of these measures were originally developed for Euclidean distance.

The first measure is the Davies-Bouldin Index (DBI) ^54^. It quantifies the compactness of a cluster by the average distance of the cells in it from the centroid, and the separation of two clusters by the distance between their centroids. Specifically, for any two clusters *C_x_* and *C_y_* with *n_x_* and *n_y_* cells and centroids 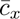 and 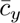, respectively, a compactness-to-separation ratio, *R_xy_*, is first computed:

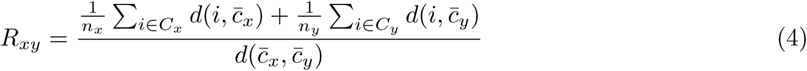

A larger value of this ratio corresponds to a worse separation of the clusters relative to their compactness. Next, for each cluster, another cluster that gives the worst compactness-to-separation ratio is identified and finally, these worst-case ratios are averaged over all clusters to form the overall DBI:

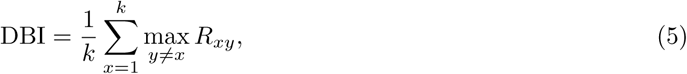

where *k* is the total number of clusters.

The second measure is the Average Silhouette Width (ASW) ^55^. For each cell *i* from a cluster *C_x_* with *n_x_* cells, the average distance between *i* and all other cells in the same cluster is first computed:

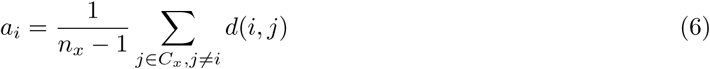

Similarly, the average distances between cell *i* and cells in each of the other *k*−1 clusters are computed, and the smallest one among them is selected:

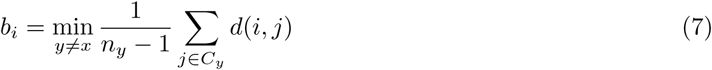

Finally, the difference between the inter-cluster distance and the intra-cluster distance is computed, normalized, and sum over all *n* cells in the data set to give the overall ASW:

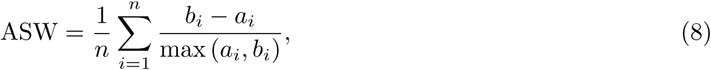

ASW has a range of −1 to 1. A larger value of ASW corresponds to a better separation of the clusters relative to their compactness.

The third measure is the Calinski-Harabasz Index (CHI) ^56^. It quantifies the compactness of a cluster by the distances of the cells from its centroid, and the separation of all clusters by the distances of each cluster’s centroid to the centroid of the whole set of cells. CHI then computes a normalized ratio between them. Specifically, denote the centroid of each cluster *C_x_* with *n_x_*cells as 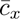 and the centroid of the whole data set with *n* cells as 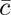, CHI is defined as:

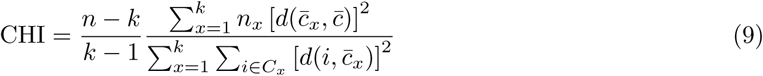

A larger value of CHI corresponds to a better separation of the clusters relative to their compactness.

#### External validation measures for evaluating agreement between cell clusters and externally obtained cell type labels

When an externally acquired reference can be used to define cell type labels of individual cells, we compared these cell type labels with the cell clusters produced as another way to evaluate the clusters. For each cluster, if ≥ 50% of its cells come from the same cell type according to the external reference, we label the whole cluster by that cell type. Otherwise, the cluster receives a special “unannotated” label. Based on this, for each cell type we defined the following:

- TP (true positive): proportion of cells annotated with this cell type that are put in a cluster also labeled by this cell type
- FP (false positive): proportion of cells not annotated with this cell type but are put in a cluster labeled by this cell type
- TN (true negative): proportion of cells not annotated with this cell type that are put in a cluster also not labeled by this cell type
- FN (false negative): proportion of cells annotated with this cell type that but are put in a cluster labeled by this cell type

With these definitions, for each cell type we further defined precision, recall, and the F1 measure:

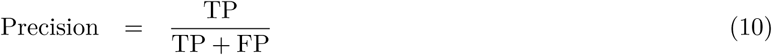

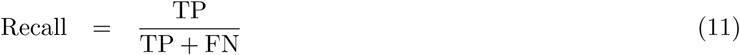

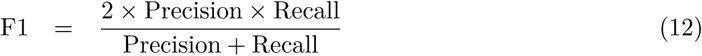

We remark that our use of these terms here is non-standard and therefore they should be interpreted with caution for the following two reasons. First, labeling each cluster by the cell type occupying ≥ 50% of its cells is just one way to enable the calculation of these performance measures rather than a “ground truth”. Second, clusters with no cell types occupying ≥ 50% of its cells receive the special “unannotated” label, which does not exist when these terms are used for evaluating performance in a standard classification setting.

The fourth measure is the Adjusted Rand Index (ARI) ^57^. The Rand Index (RI) ^58^ is the ratio of cell pairs that are consistent between the clustering results and the external reference. A cell pair is considered consistent between the two if either they are from the same cluster and are given the same cell type label by the external reference, or they are not from the same cluster and are given different cell type labels by the external reference. ARI further adjusts RI by considering the maximum achievable RI and the expected RI given the number of cells in each cluster and each cell type:

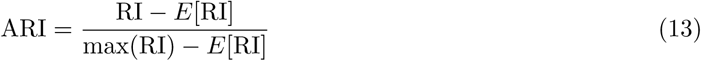

ARI ranges from −1 to 1, with a larger ARI corresponding to a better agreement between the clustering results and the external reference. A perfect agreement between the two would give an ARI of 1, while a random assignment of cells to the clusters is expected to give an ARI of 0.

The fifth measure is the Normalized Mutual Information (NMI) ^59^. For a random variable *C* with *k* possible outcomes 1*..k*, its entropy H(*C*) is defined as

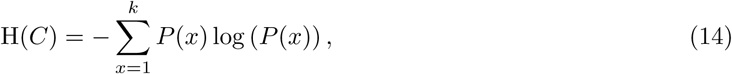

where *P* (*x*) is the probability that the variable has outcome *x*. In our current context, suppose all the cells are partitioned into *k* clusters 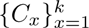 and thus each cell belongs to one of the *k* clusters (i.e., outcomes), by estimating *P* (*x*) empirically by the fraction of cells that are in cluster 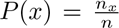 quantifies the uncertainty in guessing which cluster a cell belongs to.

Based on the entropy, the Mutual Information (MI) is defined as the amount of information that two variables share, which is quantified by the reduction in uncertainty of one variable when the other variable is known. When comparing two partition of the cells, namely a set of *k* clusters *C* and an externally obtained set of cell type labels 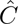 that assigns each cell to one of the 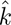 cell types (where 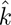 does not necessarily equal *k*), their mutual information 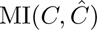 is defined as:

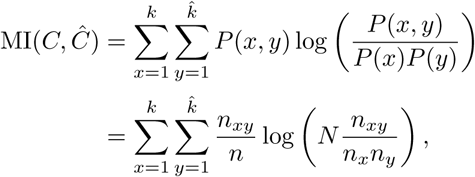

where *P* (*y*) = *n_y_/n* is the fraction of cells that belong to cell type *y* according to the external reference, and *P* (*x, y*) = *n_xy_/n* is the fraction of cells that belong to both cluster *x* and cell type *y*.

NMI further normalizes MI by dividing it by the average entropy of the two variables:

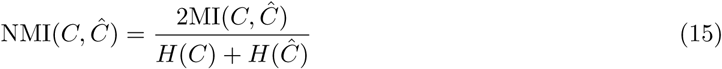

NMI has a range of 0 to 1, where a higher value indicates a better match between the clusters and the externally obtained cell type labels.

### Procedures for annotating cells in the pancreas data set with specific islet endocrine subtypes and for identifying cells with suspicious cell type annotations

We developed a procedure to annotate cells in the pancreas data set with specific islet endocrine subtypes (Figure S9). First, we used SAKURA to produce an embedding of all the cells in the pancreas data set, selected cells in clusters that express the hormone genes (Figure 3a), re-clustered them, and selected the clusters that were composed of *>* 50% of cells annotated as “islet endocrine cells” by the original authors. We then focused on cells within these clusters that were either not given a cell type annotation or given only the broad annotation of “islet endocrine cells” by the original authors.

For the originally unannotated cells, we performed doublet detection using three methods, namely DoubletDecon ^60^, DoubletFinder ^61^, and scDblFinder ^62^, using either default parameter settings or configuration recommendations given in the tutorials provided by the authors. For each method, we took its binary output to classify a cell as a doublet or not a doublet. We then combined the results of the three methods and only kept cells that were not classified as a doublet by any of the three methods.

Next, we considered three quality control (QC) metrics of each cell, namely total number of reads, total UMI count, and number of detected genes. Non-doublet cells that have quality issues typically have a low value in one of these metrics. We considered values within the lowest 30 percentile among all non-doublet cells to be potentially problematic, and kept only cells that do not have problematic values in all three metrics.

In both the filtering based on doublet detection and QC metrics, we used the conservative thresholds to make sure that all cells that were kept are in good data quality. All these cells were then temporarily given the broad label of “islet endocrine cells”.

For all high-quality cells given the broad label of “islet endocrine cells”, by our procedure or by the original authors, we further checked whether it was in a SAKURA-based cluster that expresses only one hormone gene. If that was true, we annotated the cell with the more specific islet endocrine cell subtype according to the hormone gene expressed.

For cells in these hormone-expressing clusters (including clusters that express one hormone gene or multiple hormone genes), if they were given a non-islet cell type label (e.g., “acinar cells”) by the original authors, we marked the original annotation as suspicious.

### Testing the robustness of SAKURA by providing different lists of genes of interest

We used the COVID-19 data set to test how much the performance of SAKURA is affected by the list of knowledge genes supplied. We tested 9 different gene lists, which are categorized into three groups:

- Group I: As mentioned in the main text, having *CD4* as one of the knowledge genes was not very useful in separating CD4^+^ and CD8^+^ T cells because some CD4^+^ T cells, as indicated by their surface protein levels, have low expression of *CD4* at the transcript level. Therefore, in the first group of gene lists, we focused on supplying CD8 genes as input knowledge and tested how different combinations of them affected the performance of SAKURA.
- Group II: To see whether some marker genes of CD4^+^ T cells are useful at all, we tested different combinations of their makers.
- Group III: To see how SAKURA’s performance is affected by genes unrelated to the main goal of separating CD4^+^ and CD8^+^ T cells, we added some irrelevant genes.

Together, these three groups cover different scenarios that have incomplete or noisy knowledge supplied as input to SAKURA.

Specifically, the 9 gene lists were defined as follows:

- **Group I SUM(CD8):** the knowledge input specifies the sum of the expression of the *CD8A* and *CD8B* genes as the regression target.
- **Group I CD8A, CD8B:** the two genes that encode the alpha and beta subunits of the CD8 glycoprotein were supplied as the knowledge genes, which serve as separate regression targets. This represented another way to take the two genes as knowledge input.
- **Group I CD8A:** only the *CD8A* gene was supplied as knowledge input, to see how missing *CD8B* in the knowledge input would affect SAKURA’s performance. Since *CD8A* is more consistently and highly expressed across CD8^+^ T cells compared to *CD8B* in our observations and previous reports ^63^, we expected that *CD8A* would be the more reliable marker of these cells in single-cell analysis.
- **Group I CD8B:** only the *CD8B* gene was supplied as knowledge input, to see how missing *CD8A* in the knowledge input would affect SAKURA’s performance.
- **Group II CD4:** only the *CD4* gene was supplied as knowledge input. Again, this setting was expected to be not very useful for separating CD4^+^ and CD8^+^ T cells.
- **Group II CCR7_IL7R:** as alternative markers of CD4^+^ T cells, *CCR7* (C-C chemokine receptor type 7) and *IL7R* (interleukin-7 receptor) were supplied as knowledge input.
- **Group III SUM(CD8)-n-IFNG:** The knowledge supplied included two inputs, namely the *IFNG* gene and the sum of *CD8A* and *CD8B*. *IFNG* encodes interferon-gamma (IFN-*γ*), which is a functional marker for identifying effector T cells but is expected to be irrelevant to separating CD4^+^ and CD8^+^ T cells. The “-n-” part indicates that the gene after it (*IFNG*) is a noisy knowledge input.
- **Group III SUM(CD8)-n-LYZ:** The knowledge supplied included two inputs, namely the *LYZ* gene and the sum of *CD8A* and *CD8B*. *LYZ* encodes lysozyme, which is expressed in various activated immune cells but is expected to be irrelevant to separating CD4^+^ and CD8^+^ T cells.
- **Group III SUM(CD8)-n-FN1:** The knowledge supplied included two inputs, namely the *FN1* gene and the sum of *CD8A* and *CD8B*. *FN1* encodes fibronectin, which is involved in the regulation of some immune responses but is expected to be irrelevant to separating CD4^+^ and CD8^+^ T cells.

## Declarations

### Ethics approval and consent to participate

Not applicable

### Consent for publication

Not applicable

### Data availability

All data used in this study are publicly available. The Human Fetal Cell Atlas scRNA-seq data are available at https://descartes.brotmanbaty.org/bbi/human-gene-expression-during-development/ ^64^. The COVID-19 CITE-seq data from the COMBAT consortium are available at https://zenodo.org/records/6120249^65^. The snRNA-seq data from aging brain are available at https://cellxgene.cziscience.com/collections/31937775-0602-4e52-a799-b6acdd2bac2e ^66^. The SAKURA method is implemented with Python and is freely available at its Github repository https://github.com/Yip-Lab/SAKURA ^67^ and Zenodo https://zenodo.org/records/15801301^68^ under the MIT license.

### Authors’ contributions

KYY conceived the study. ZZ, JC, HW, KYL, and KYY designed the methods. ZZ, JC, HW, and KYL implemented the methods, collected and processed the data, and performed the data analysis. ZZ, JC, HW, KYL, PA, PI-A, and KYY interpreted the results. ZZ, JC, HW, KYL, PI-A, and KYY wrote the manuscript. All authors read and approved the final version of the manuscript.

