## Supplementary Materials for "A knowledge-guided approach to recovering important rare signals from high-dimensional single-cell data"

### Supplementary information

#### Supplementary figures

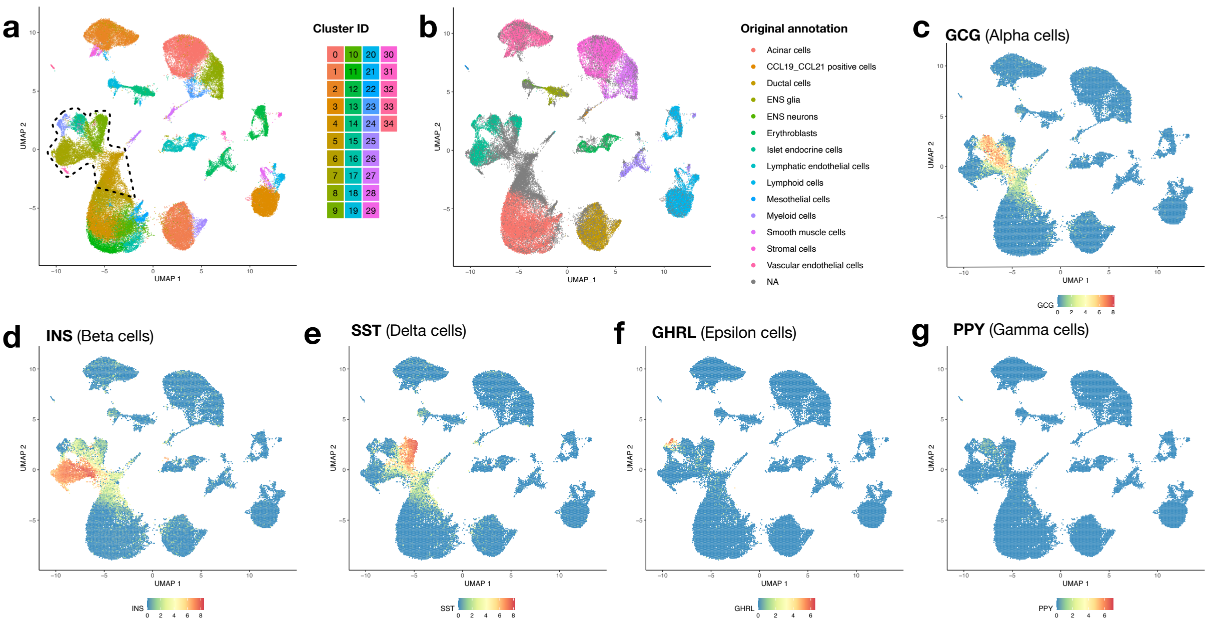


**Supplementary Figure S1: Unsupervised analysis of the pancreas data set. a,** Cell clusters formed by the unsupervised pipeline (PCA followed by clustering; Methods). Cells coming from each cluster are shown in the same color. Clusters that contain most of the cells expressing the five hormone genes are marked by the black contour line. **b,** UMAP visualization of the pancreas data set with the original cell type annotations. **c-g,** Expression levels of the five hormone genes in all cells, namely *GCG* (c), *INS* (d), *SST* (e), *GHRL* (f), and *PPY* (g).

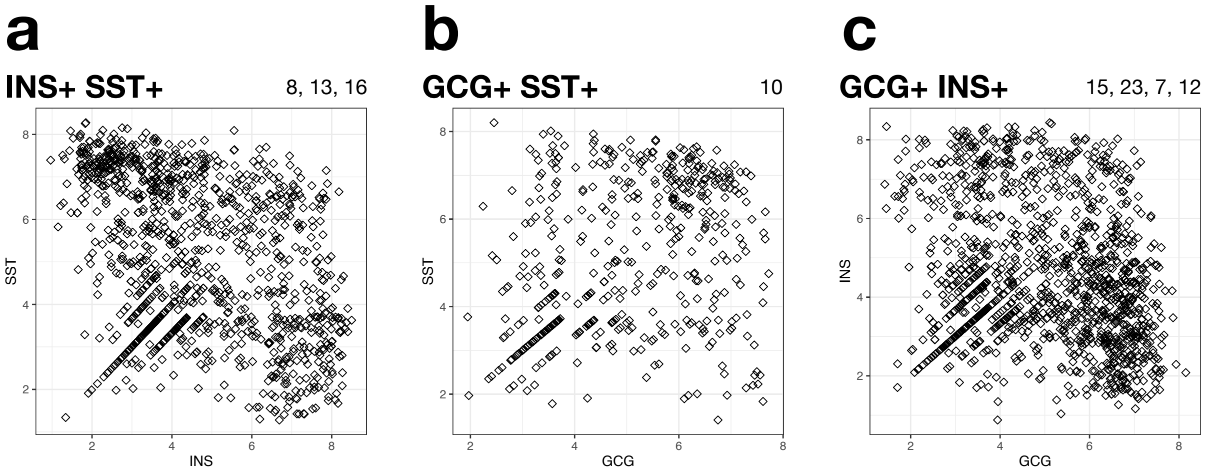


**Supplementary Figure S2: Cells in clusters that express multiple hormone genes. a-c,** Expression levels of two different hormone genes in each cell that comes from clusters expressing multiple hormone genes, namely *INS* and *SST* (a), *GCG* and *SST* (b), and *GCG* and *INS* (c). The numbers above the scatterplots are identifiers of the clusters from which the cells are extracted.

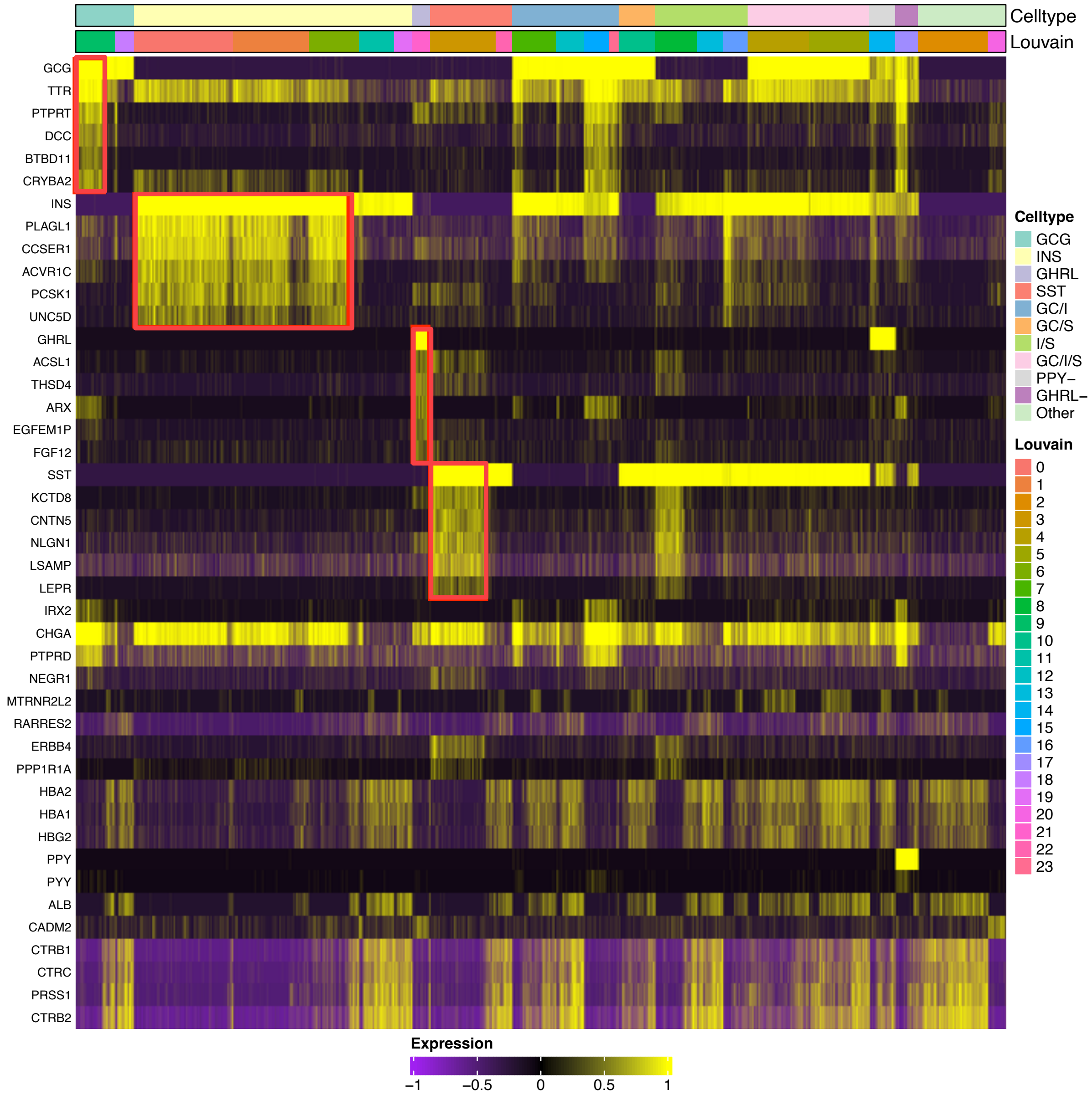


**Supplementary Figure S3: Differentially expressed genes between the hormone gene expressing cell groups.** The heatmap shows relative expression levels of genes differentially expressed between cells in groups that express different combinations of hormone genes. All genes shown have a log fold change > 0.25, and for each hormone gene combination, up to 6 genes with the strongest fold changes are shown. Each row corresponds to one of these genes and each column corresponds to a cell. Cells from the same Louvain clusters are placed next to each other, and clusters that express the same set of hormone genes, which form a group and are given the same cell type label, are also placed next to each other. The red boxes show several sets of genes that correlate with hormone genes in cluster-specific manners.

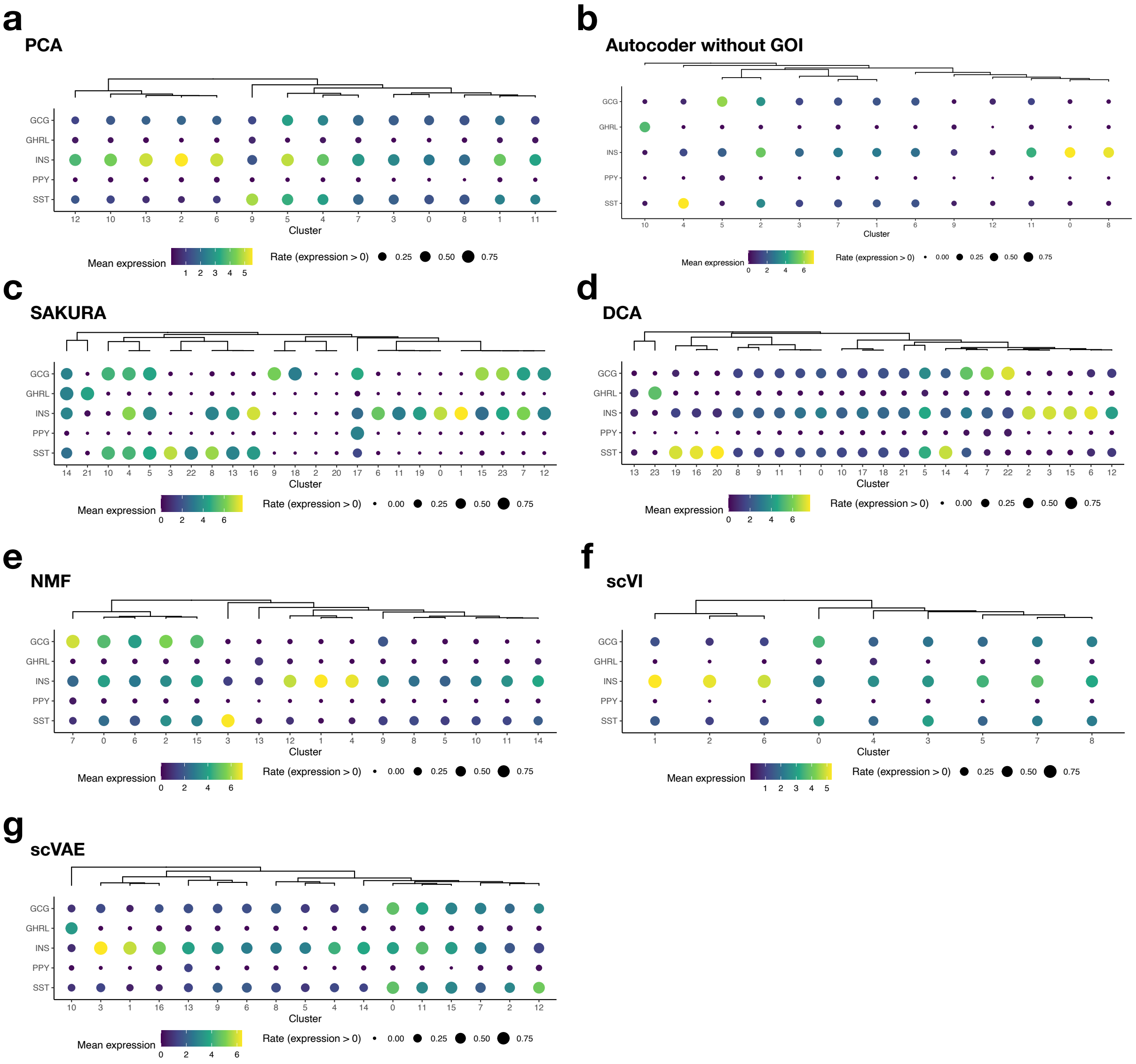


**Supplementary Figure S4: Expression of the five hormone genes in the cell clusters based on cell embeddings produced by different methods. a-g,** Expression of the five genes based on cell embeddings produced by PCA (a), Autoencoder without GOI (b) SAKURA (c), DCA (d), NMF (e), scVI (f), and scVAE (g). In all cases, the cells involved are those within the contour line shown in Figure S1a.

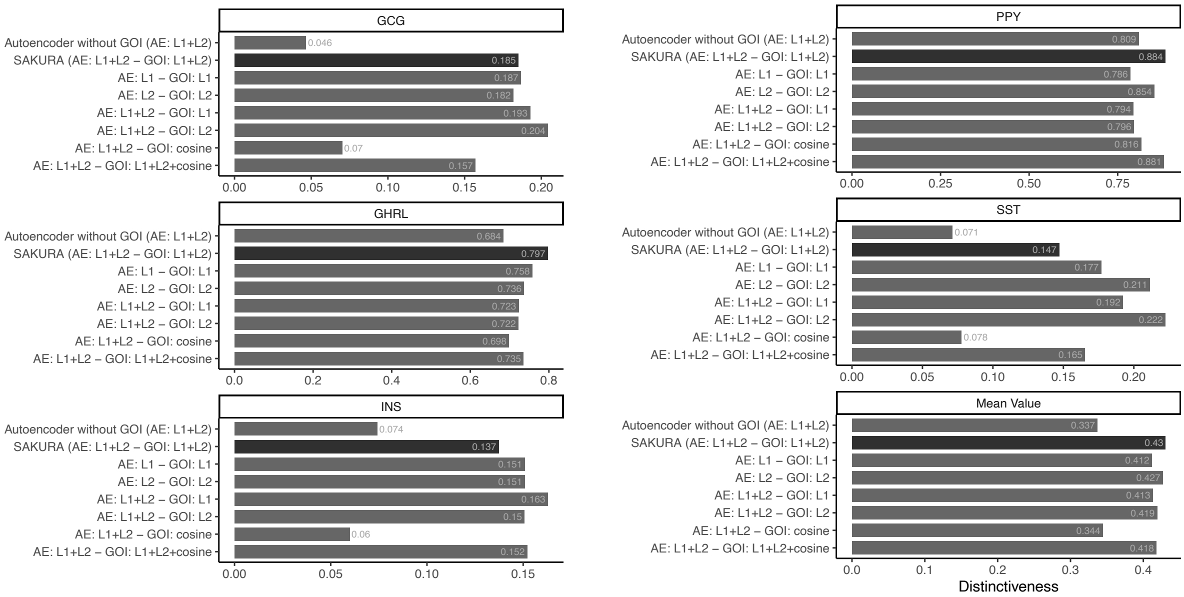


**Supplementary Figure S5: Distinctiveness of expression of the hormone genes in cell clusters produced from SAKURAs embeddings.** Average distinctiveness across all clusters of expression of the hormone genes the produced from SAKURA's autoencoder backbone (Autoencoder without GOI) and different loss term configurations of the autoencoder backbone (AE) and the GOI module. Mean distinctiveness values of the hormone genes for each setting are also presented. In all cases, the cells involved are those within the contour line shown in Figure S1a.

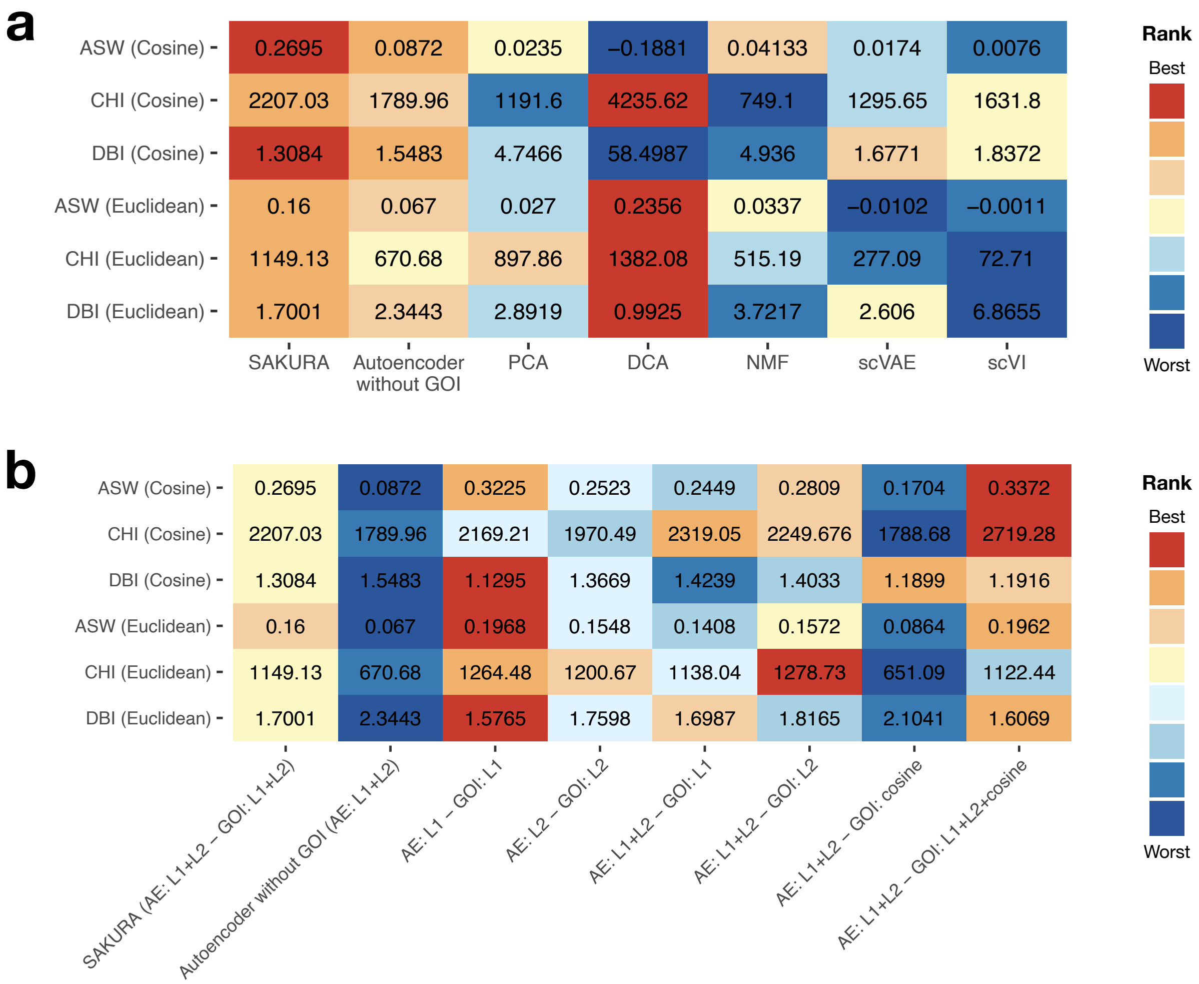


**Supplementary Figure S6: Internal validation of different cell embedding methods applied to the pancreas data set.** Three internal validation measures are used to evaluate the embeddings produced by **a,** SAKURA, SAKURA's autoencoder backbone and five unsupervised methods (columns) and **b,** SAKURA with different loss term configurations of the autoencoder backbone(AE) and the GOI module (columns). ASW (larger is better), CHI (larger is better), and DBI (smaller is better), based on clusters produced from k-nearest neighbor graphs constructed according to cosine distance (first three rows) or Euclidean distance (last three rows). In each row, ranking of the methods is indicated by the colors. In all cases, the cells involved are those within the contour line shown in Figure S1a.

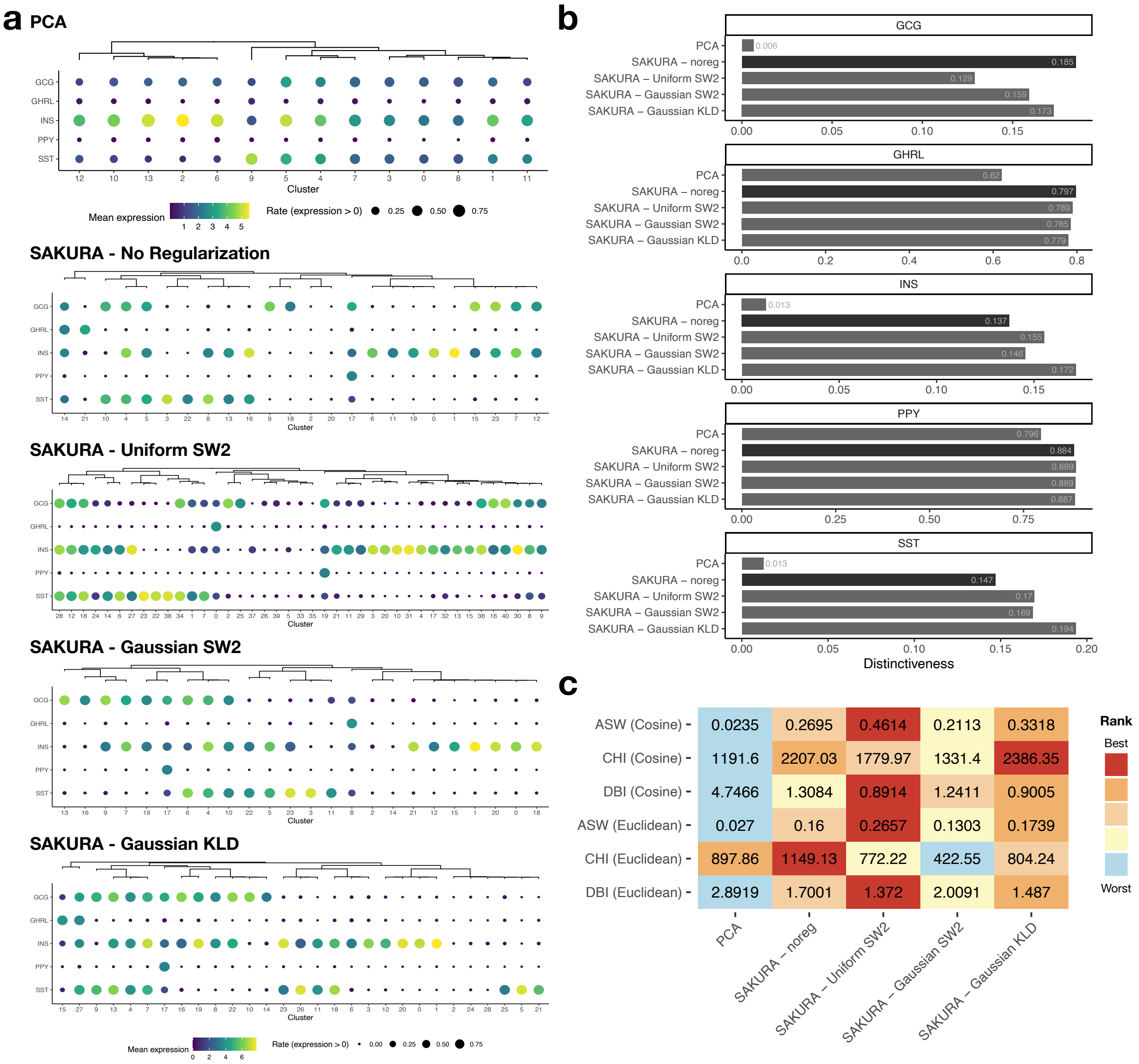


**Supplementary Figure S7: Comparative analysis of SAKURA regularization configurations for Islet cell cluster identification.** **a,** Expression of the five hormone genes in the clusters of the extracted cells identified by the unsupervised pipeline and clusters from SAKURA's embeddings with different regularization configurations. In Panels a, a complete-link hierarchical clustering of the cell clusters was performed based on cosine similarity of the detection rates of the five hormone genes. **b,** Distinctiveness of the expression of the hormone genes in cell clusters produced from the unsupervised pipeline and SAKURA's embeddings with different regularization configurations. For each gene, the distinctiveness shown is the average across all clusters. **c,** Internal validation of different cell embeddings from the unsupervised pipeline and SAKURA with different regularization configurations.ASW (larger is better), CHI (larger is better), and DBI (smaller is better), based on clusters produced from k-nearest neighbor graphs constructed according to cosine distance (first three rows) or Euclidean distance (last three rows). In each row, ranking of the methods is indicated by the colors. In all cases, the cells involved are those within the contour line shown in Figure S1a.

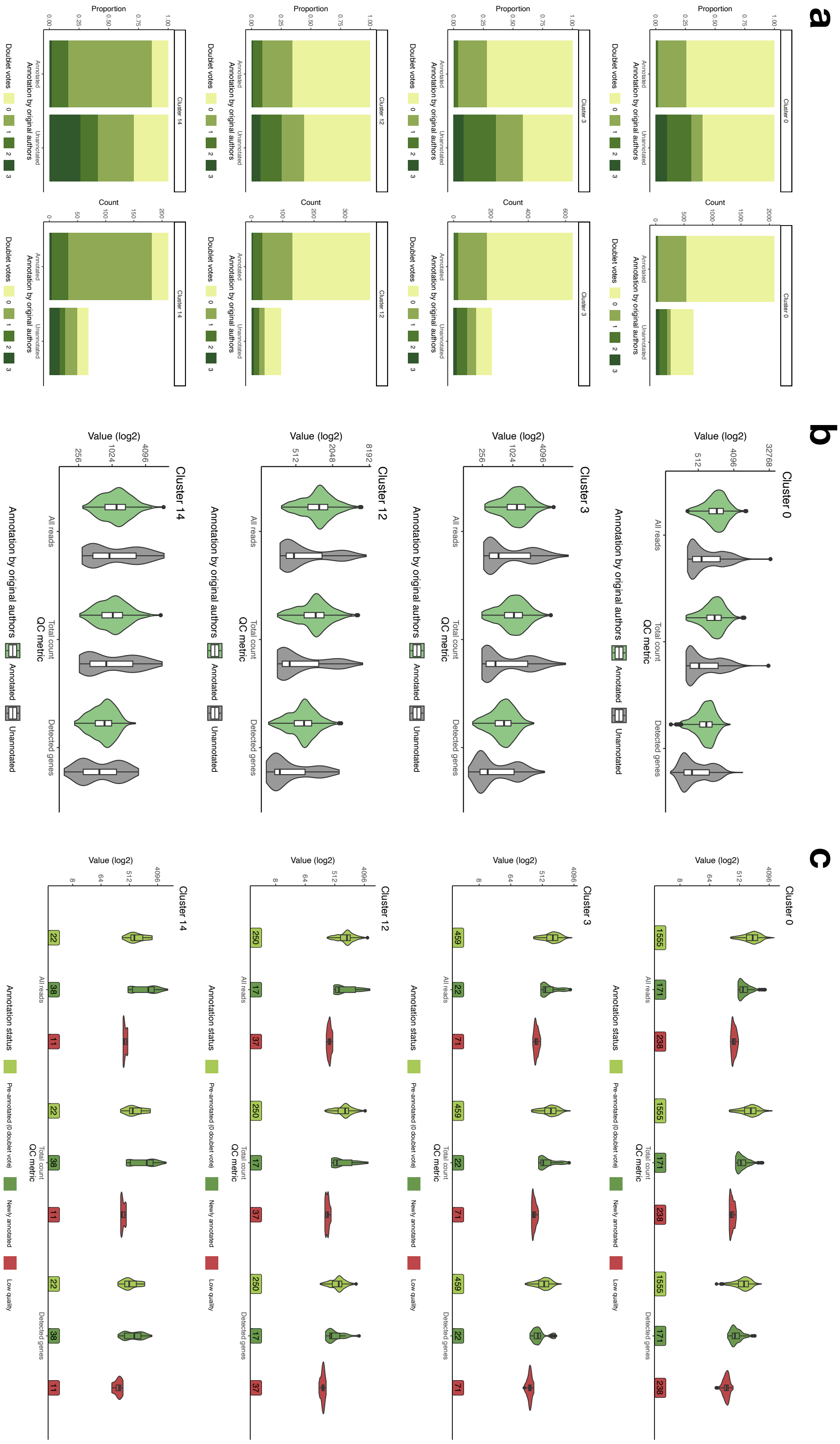


**Supplementary Figure S8:** (Figure caption on the next page.)

**Supplementary Figure S8: Doublet votes, QC metrics, and reassignment recommendations.** **a,** Comparing the doublet votes of cells annotated and unannotated by the original authors in clusters dominated by pre-annotated pancreatic islet cells. The two columns show the proportions and absolute counts of cells, respectively. **b,** Comparing the QC metrics of cells annotated and unannotated by the original authors in these clusters. **c,** Comparing QC metrics of cells with different cell type label reassignment recommendations in these clusters. Cells in the “Pre-annotated (0 doublet vote)” group were annotated by the original author and not reported as doublets by any of the three doublet detection methods we used. Cells in the “Newly annotated” group are the originally unannotated cells in our selection that passed our doublet and quality requirements. Cells in the “Low quality” group are the originally unannotated cells in our selection that did not pass our doublet or quality requirements.

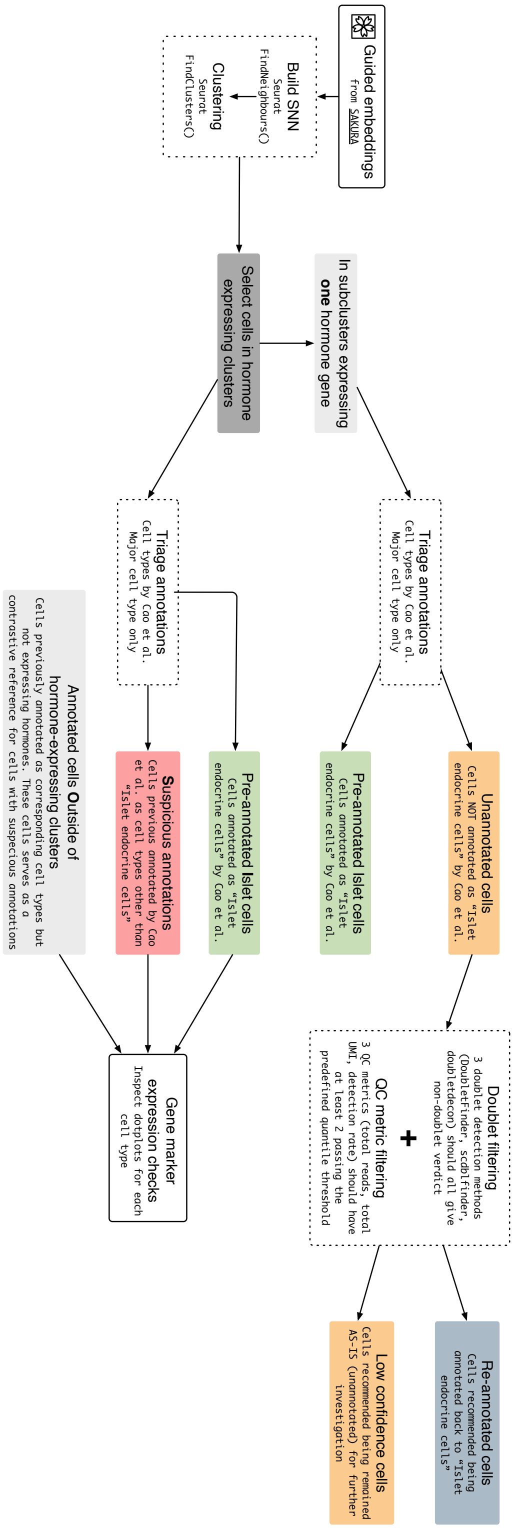


**Supplementary Figure S9: Workflows for improving cell type annotations.** Two workflows were developed, namely one for identifying cells of specific islet endocrine cell subtypes, and one for identifying cells with suspicious cell type labels given by the original authors.

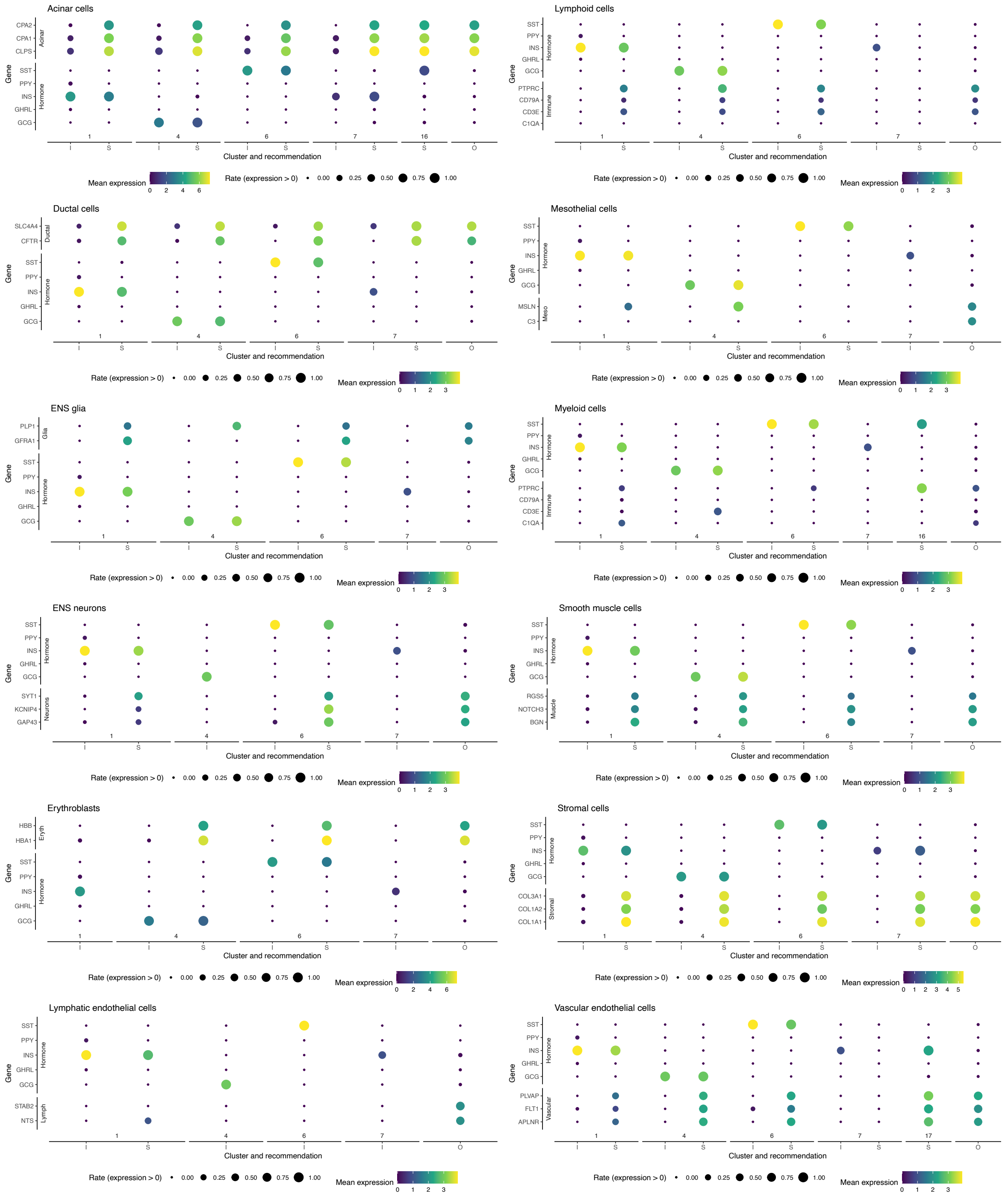


**Supplementary Figure S10: Expression of marker genes of different cell types and the hormone genes in the cells identified by SAKURA as requiring re-checking.** Each sub-figure is about one non-islet cell type in the pancreas. Expression of marker genes of the cell type and the hormone genes in the hormone-expressing cell clusters identified by SAKURA is shown for different groups of cells. I: cells annotated by the original authors as islet endocrine cells. S: cells annotated as that non-islet cell type by the original authors that are marked as suspicious by SAKURA. O: cells annotated as that non-islet cell type by the original authors that are outside the hormone-expressing clusters produced by SAKURA, which serve as controls of cells of that non-islet cell type.

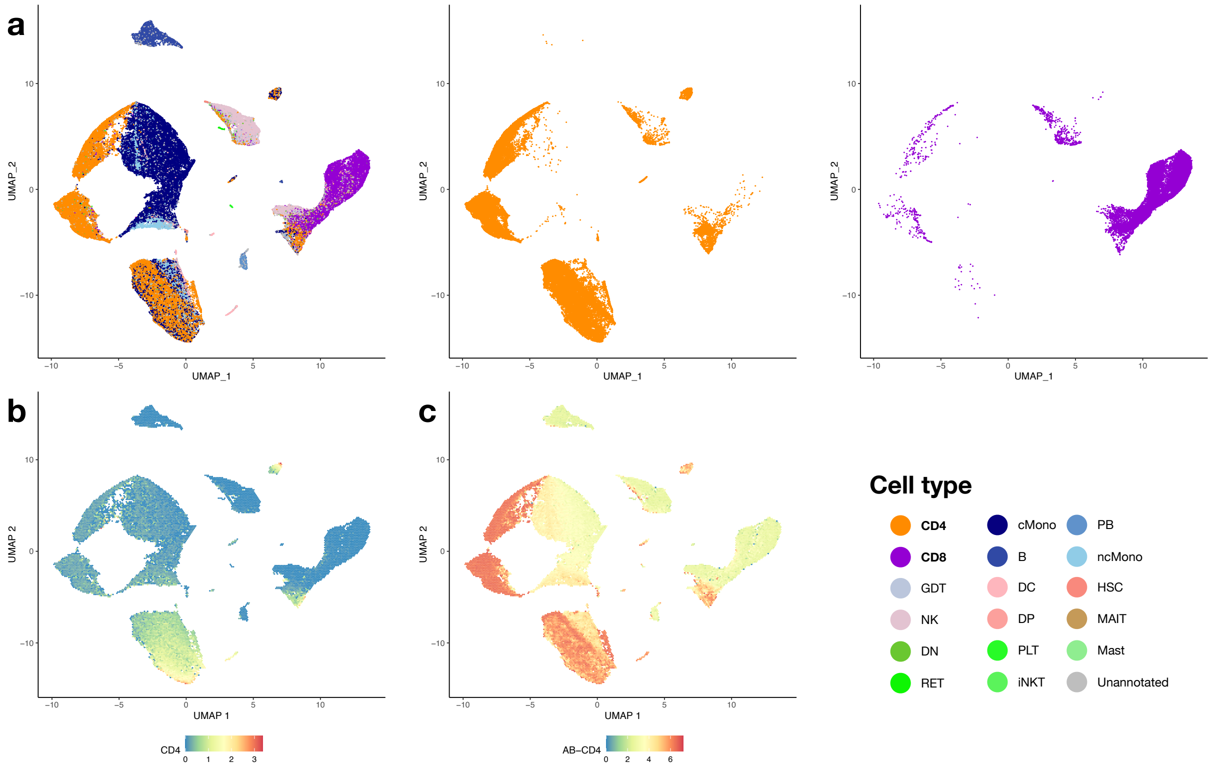


**Supplementary Figure S11: Guiding the analysis of the COVID-19 data by supplying *CD4* and *CD8A* + *CD8B* as knowledge input. a,** UMAP visualization of the cells based on SAKURA's embedding, showing all cell types, only CD4$^+$ T cells, or only CD8$^+$ T cells. **b,** Expression of *CD4* at the transcript level. **c,** Expression of CD4 at the cell surface protein level.

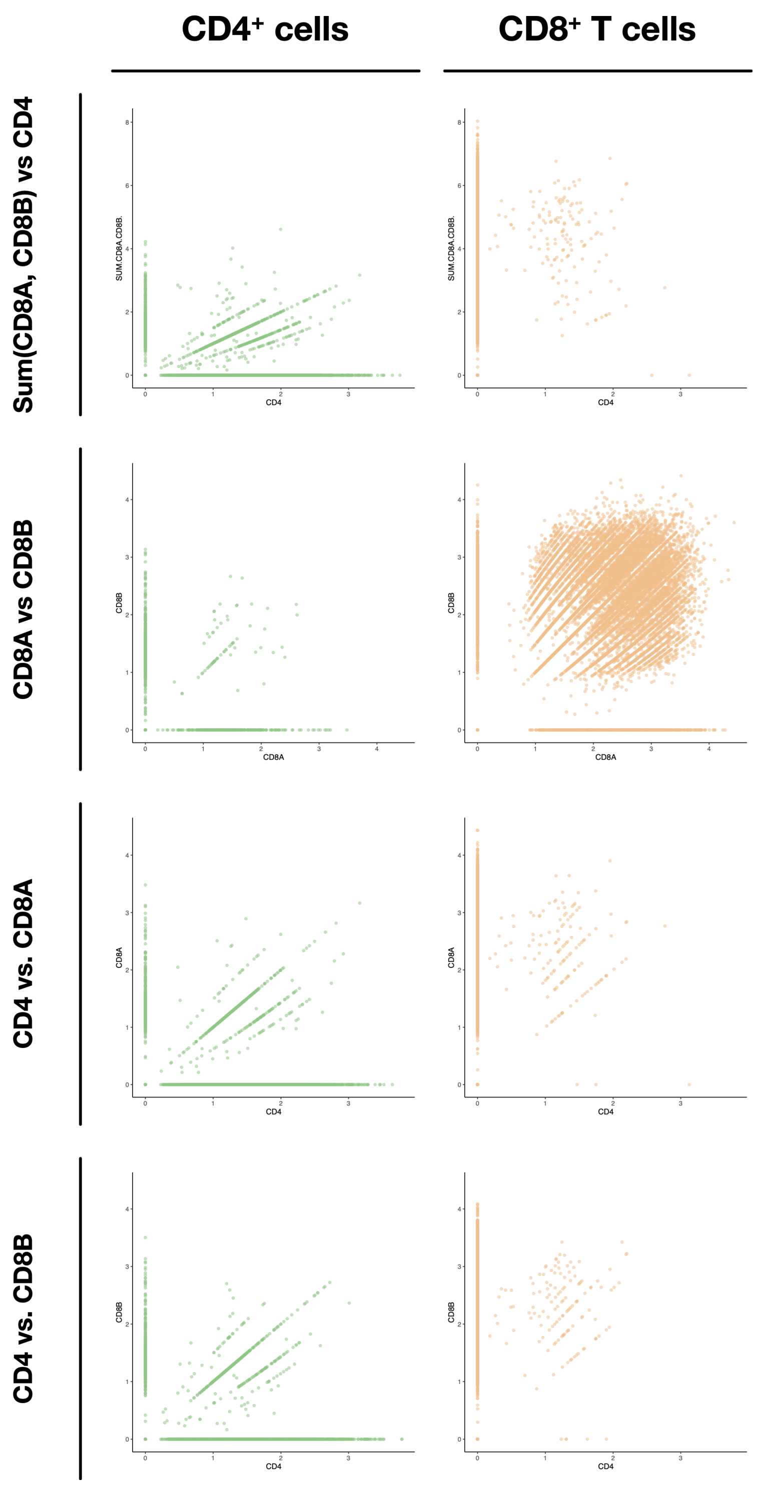


**Supplementary Figure S12: Expression of marker genes in annotated CD4$^+$ and CD8$^+$ cells.** The rows show different combinations of marker genes and their corresponding expression levels in CD4$^+$ cells (first column) and CD8$^+$ cells (second column) annotated by the original authors using both transcript and cell surface protein information.

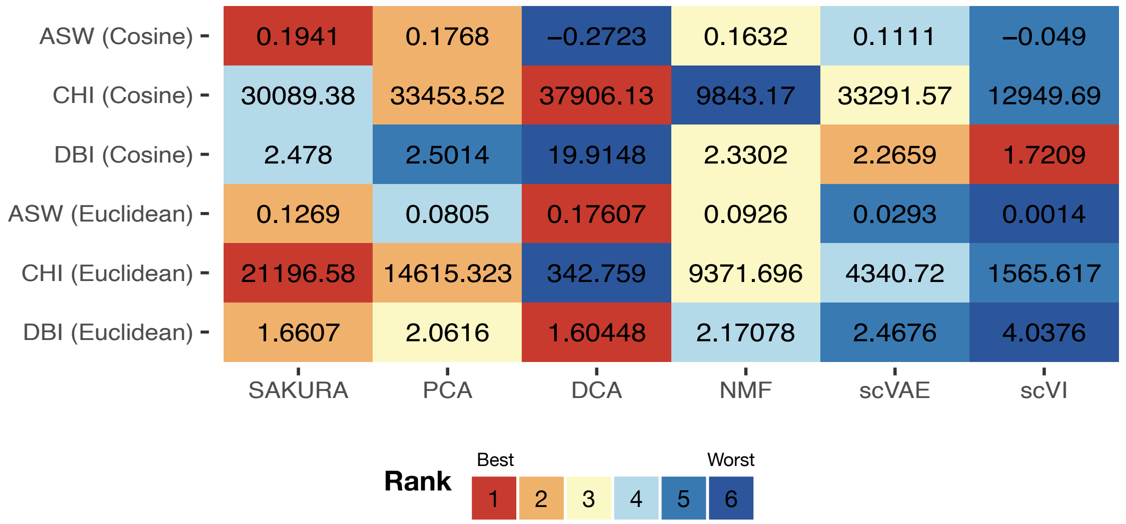


**Supplementary Figure S13: Internal validation of different cell embedding methods applied to the COVID-19 data set.** The embeddings produced by SAKURA and five unsupervised methods (columns) are evaluated based on three internal validation measures, ASW (larger is better), CHI (larger is better), and DBI (smaller is better), based on clusters produced from k-nearest neighbor graphs constructed according to cosine distance (first three rows) or Euclidean distance (last three rows). In each row, ranking of the methods is indicated by the colors.

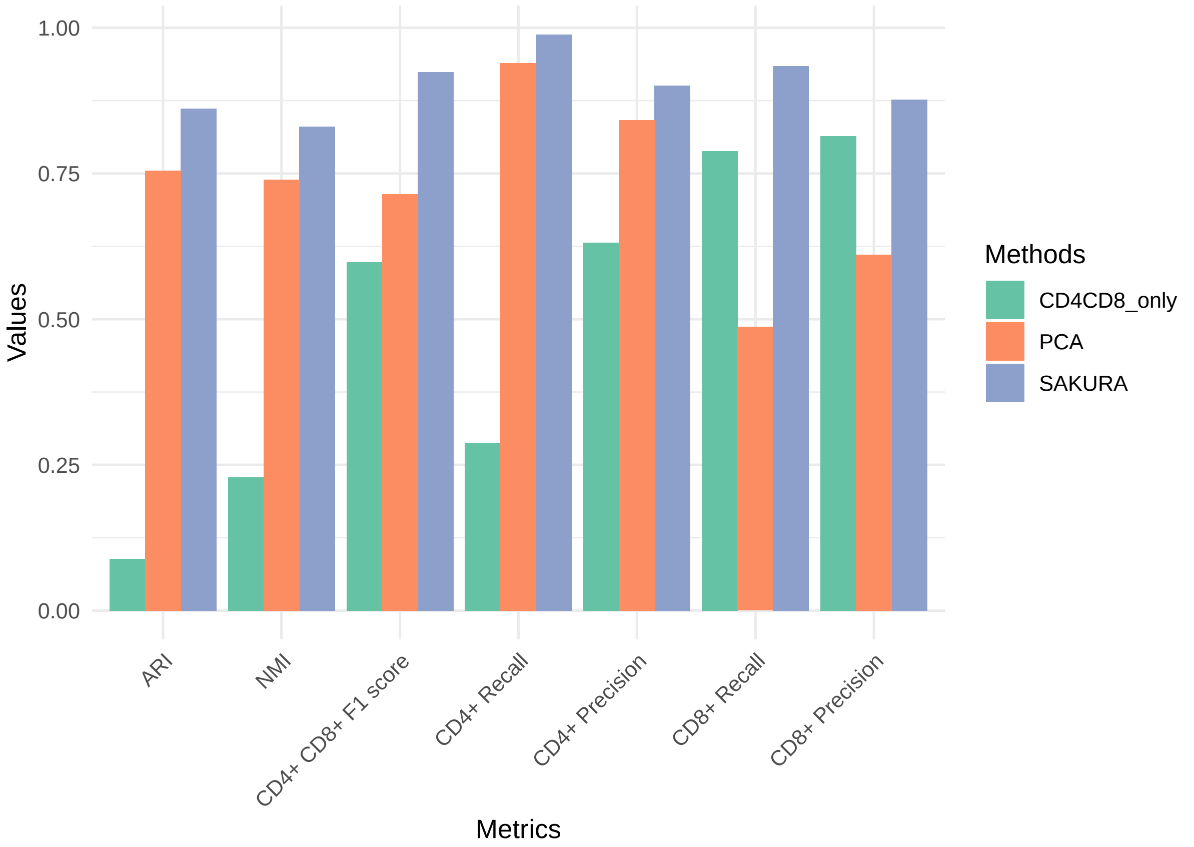


**Supplementary Figure S14: Comparing the clustering results based on the embeddings produced by either SAKURA, the standard unsupervised pipeline, or marker genes only.** Each bar group compares the clusters produced from the three embeddings based on one external evaluation measure. "CD4$^+$ CD8$^+$ F1 score" is the mean of the CD4$^+$ F1 score and the CD8$^+$ F1 score.

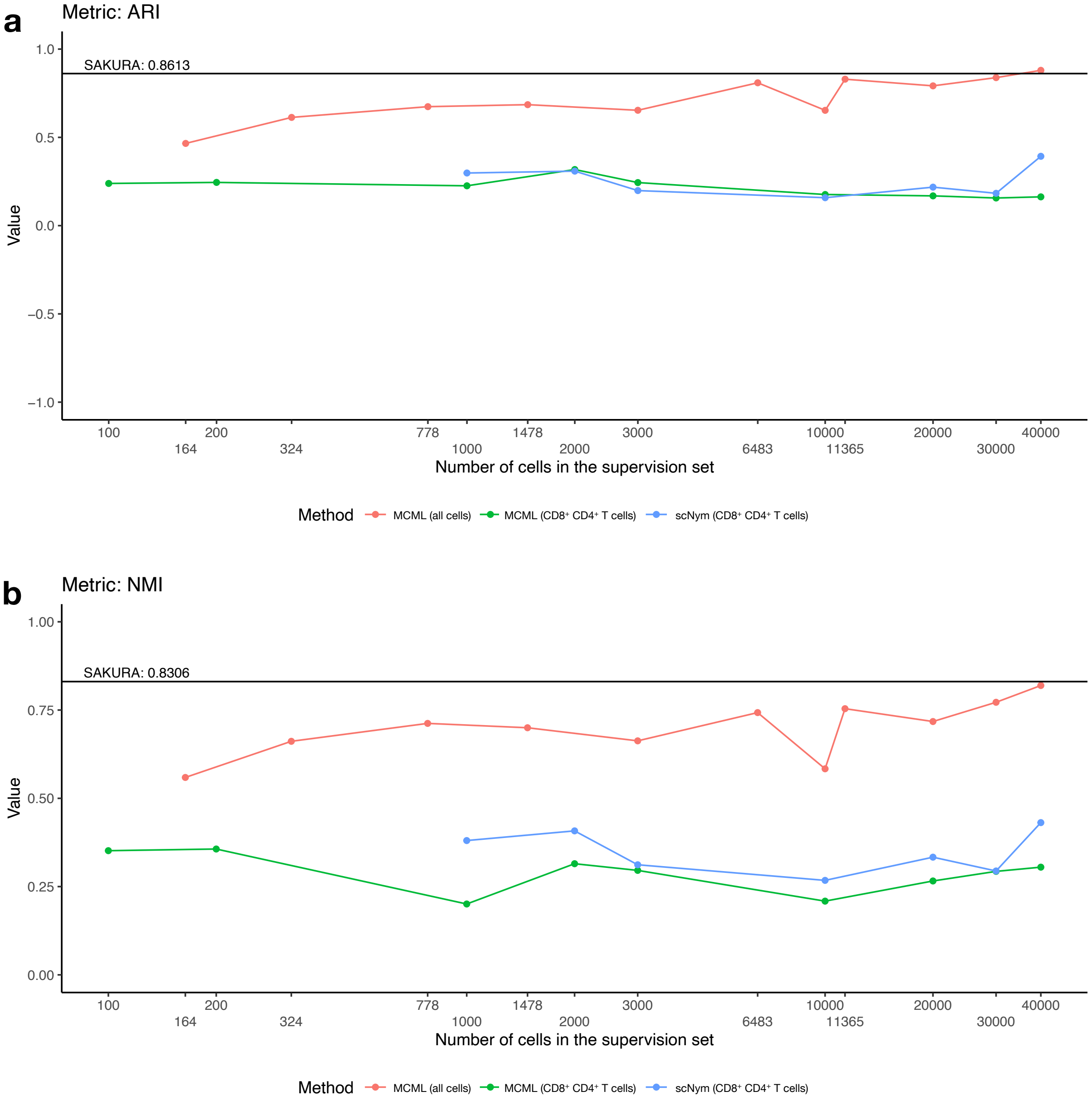


**Supplementary Figure S15: Performance of the embeddings produced by SAKURA and two semi-supervised methods that take example cells as external knowledge input in separating CD4$^+$ and CD8$^+$ cells a-b,** Performance is quantified by ARI (a) or NMI (b). The horizontal black line shows SAKURA's performance, while the three colored lines show performance of the two methods supplied with different numbers of cells as knowledge input, for all cell types or only CD4$^+$ and CD8$^+$ cells.

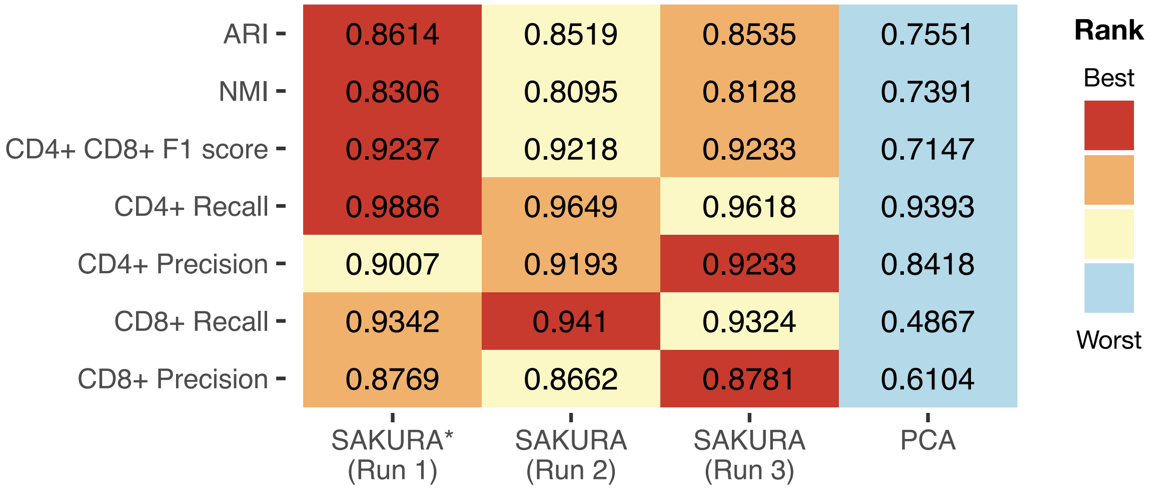


**Supplementary Figure S16: Performance of the embeddings produced by SAKURA and the unsupervised pipeline.** Performance of the embeddings produced by three runs of SAKURA and the standard unsupervised pipeline in separating $CD4^+$ and $CD8^+$ cells. Each row shows the actual performance scores and corresponding ranks based on one external evaluation measure.

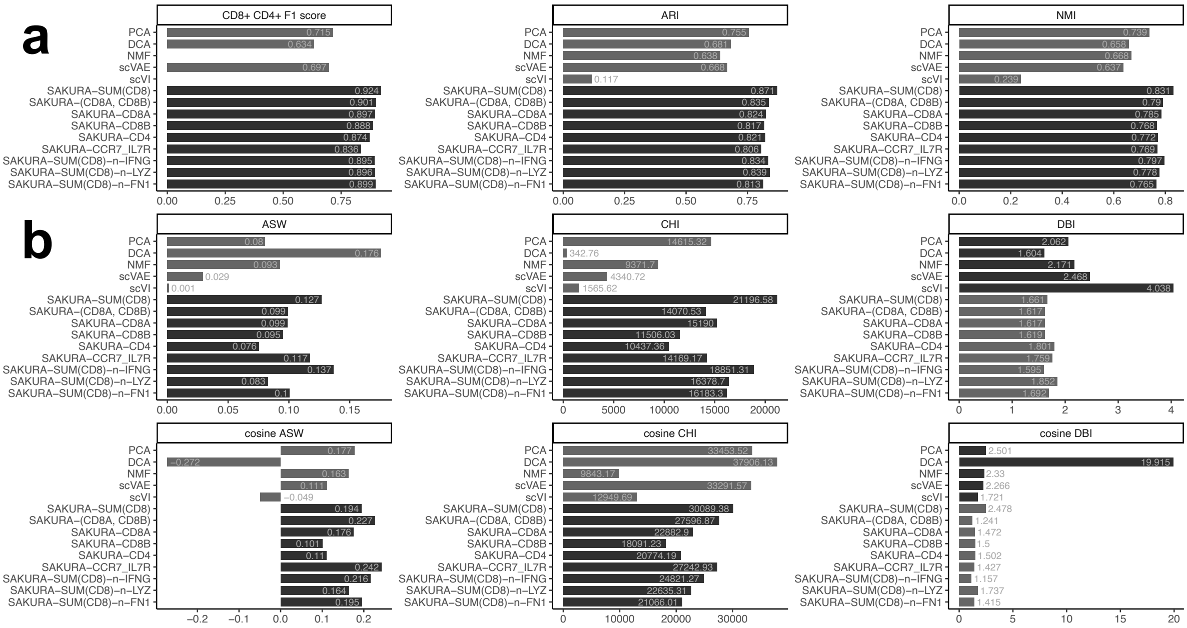


**Supplementary Figure S17: Performance of the embeddings produced by SAKURA when given different lists of genes as knowledge input. a,** Performance of the embeddings produced by SAKURA and five unsupervised methods in separating CD4$^+$ and CD8$^+$ cells, as quantified by three external validation measures, F1 score, ARI, and NMI. **b,** Data distortion assessments of the embeddings produced by SAKURA and five unsupervised methods, as quantified by three internal validation measures, ASW (larger is better), CHI (larger is better), and DBI (smaller is better), based on clusters produced from k-nearest neighbor graphs constructed according to Euclidean distance (first three rows) or cosine distance (last three rows) in the embedding. In both panels, the external knowledge provided to SAKURA is shown in the suffix, where "SUM(CD8)" means setting the sum of *CD8A* and *CD8B* transcript levels as regression target, while "-n-" indicates the gene after it is a noisy input irrelevant to CD4$^+$ and CD8$^+$ cells.

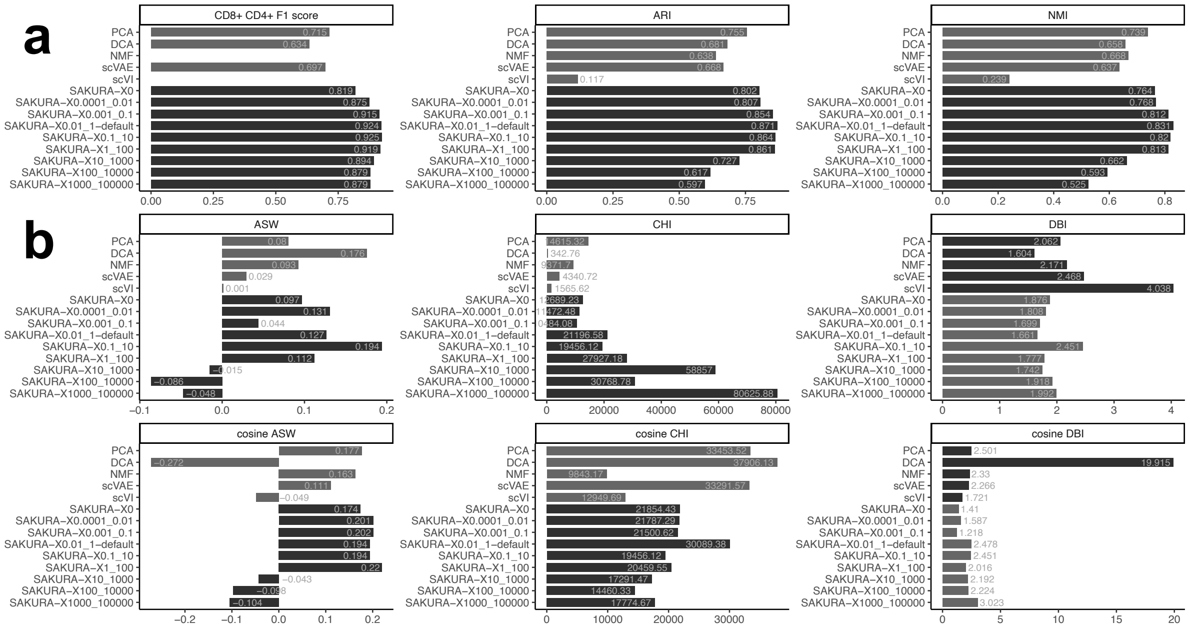


**Supplementary Figure S18: Performance of the embeddings produced by SAKURA with different supervision intensity settings. a,** Performance of the embeddings produced by SAKURA and five unsupervised methods in separating $CD4^+$ and $CD8^+$ cells, as quantified by three external validation measures, F1 score, ARI, and NMI. **b,** Data distortion assessments of the embeddings produced by SAKURA and five unsupervised methods, as quantified by three internal validation measures, ASW (larger is better), CHI (larger is better), and DBI (smaller is better), based on clusters produced from k-nearest neighbor graphs constructed according to Euclidean distance (first three rows) or cosine distance (last three rows) in the embedding. In both panels, supervision intensity settings of the knowledge input are shown in the suffix, where the number after "X" is the intensity step size and the number after "_" is the maximum intensity.

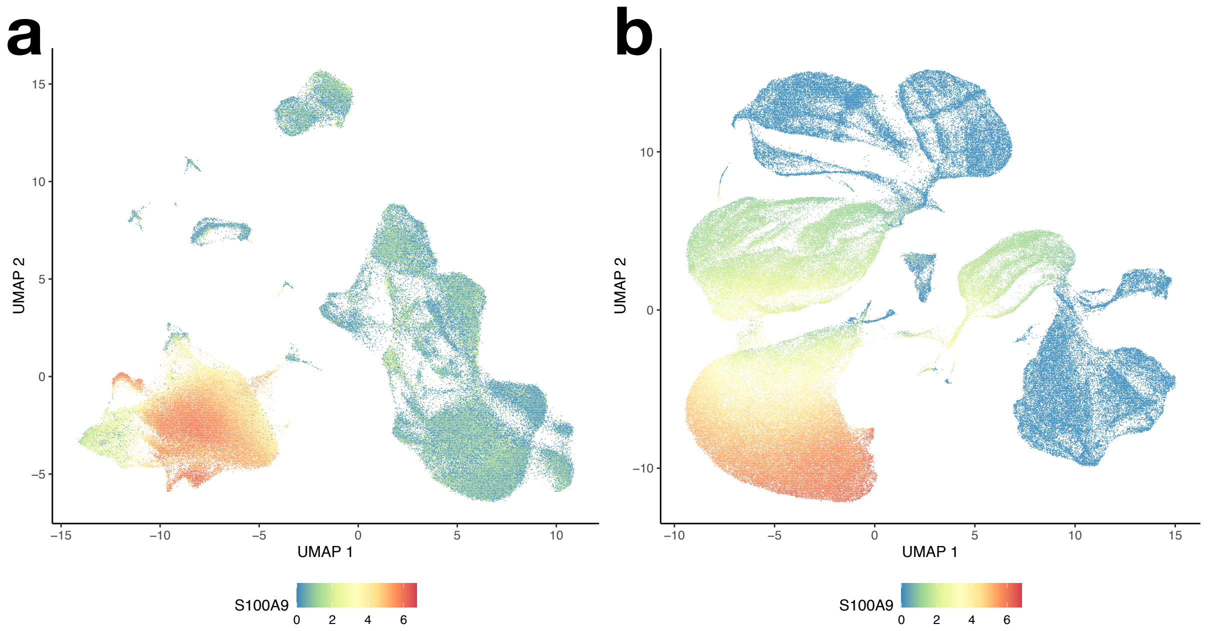


**Supplementary Figure S19: UMAP plot of cells based on different embedding pipelines. a,** UMAP plot of cells based on the embedding produced by the unsupervised pipeline applied to transcript data only. **b,** UMAP plot of cells based on the embedding produced by the SAKURA pipeline applied to transcript data only. In both panels, cells are colored by the expression level of S100A9.

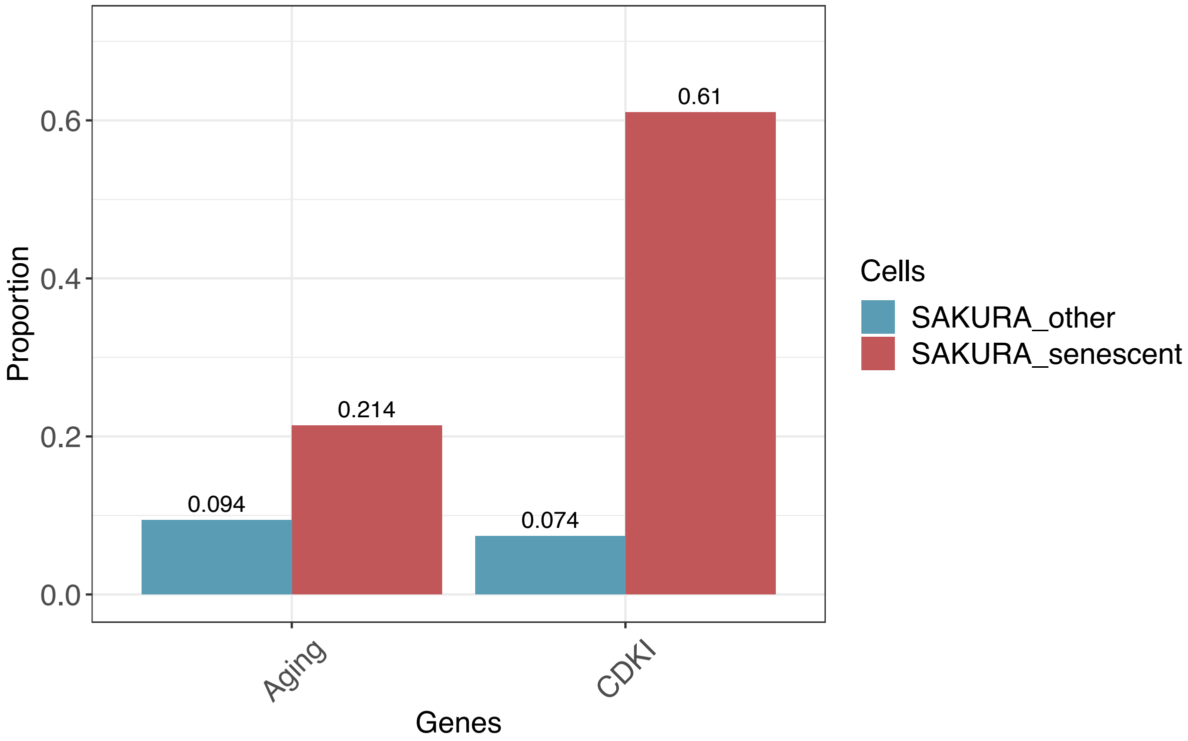


**Supplementary Figure S20:** Proportion of cells that have top 10% aging and CDKI scores in the potential senescent cells identified using SAKURA (Cluster 10, "SAKURA_senescent") and the other cells ("SAKURA_other").

#### Supplementary files

Supplementary File S1: Cells in the pancreas data set that are annotated with specific islet subtypes by SAKURA.

Supplementary File S2: Cells in the pancreas data set whose original cell type annotations are identified by SAKURA as suspicious.
